# The MLLT3 YEATS domain is a dual reader of histone marks (H3K9/18/27ac/cr) and ncRNA (7SK), linking epigenetic and RNA signaling to regulate hematopoiesis

**DOI:** 10.1101/2025.09.01.673558

**Authors:** Adam Boulton, Ashish Kabra, Nicholas Achille, Emmalee R. Adelman, Dimitrios G. Anastasakis, Felipe Beckedorff, Pradeepkumar Cingaram, Ying Zhang, Benjamin Leach, Ramin Shiekhattar, Akihiko Yokoyama, Markus Hafner, Maria Figueroa, Nancy Zeleznik-Le, John Bushweller

## Abstract

The protein MLLT3 (AF9) is a critical regulator of hematopoiesis. The N-terminal YEATS domain of MLLT3 is an epigenetic reader that binds to acetylated as well as crotonylated lysine. Using PAR-CLIP, biochemical assays, and NMR based mapping of binding, we demonstrate that the YEATS domain of MLLT3 binds to a specific stem-loop region of the noncoding RNA 7SK. 7SK is a noncoding RNA with a well-documented function in transcriptional elongation. We developed point mutations in the YEATS domain that disrupt RNA binding while having no effect on binding of acetylated histone peptides to probe the specific role of RNA binding in MLLT3 function. Our results show loss of RNA binding by MLLT3 skews hematopoietic differentiation away from the myeloid lineage and toward the lymphoid lineage and has substantial effects on gene expression, confirming the essential nature of MLLT3-RNA binding for function.

## Introduction

AF9 (MLLT3) is a critical regulator of hematopoiesis. Enver and co-workers first showed that MLLT3 is critical for the development of the erythroid/megakaryocyte lineage ^1^. More recently, Mikkola and co-workers have clearly delineated a critical role for MLLT3 in maintaining the hematopoietic stem or progenitor cell (HSPC) population ^2^. MLLT3 was localized to active promoters, enhanced levels of H3K79 methylation, and maintained a gene expression program essential for HSPCs. Daley and co-workers applied a network biology approach to identify key regulators of critical hubs for HSPCs ^3^, resulting in the identification of MLLT3 as a key regulator. Interestingly, loss of the homolog MLLT1 did not impact hematopoietic stem cell function, but has been shown to be essential for MLL fusion leukemia ^4,5^.

AF9 (MLLT3) and ENL (MLLT1) are members of the YEATS family, named after the five proteins (Yaf9, ENL, AF9, Taf14, Sas5) harboring a conserved domain termed the YEATS domain ^6^. In humans, there are four proteins with a YEATS domain: YEATS4 (GAS41), MLLT3 (AF9), MLLT1 (ENL), and YEATS2. The YEATS domain is an epigenetic reader that binds to acetylated as well as crotonylated lysine, unlike the bromodomain which can only bind to acetylated lysine. Structure determination of the AF9 (MLLT3) YEATS domain bound to acetylated and crotonylated lysine peptides ^7,8^ showed that the YEATS domain has an “end-open” binding site unlike the “side-open” site seen in bromodomains which makes it possible for the YEATS domain to accommodate the larger, more sterically demanding crotonyl modification. Structures of the YEATS domains of Taf14 and YEATS2 have also been determined ^9–11^.

Work from different laboratories, including ours, has revealed roles of MLLT3 and MLLT1 (MLLT3/1) in at least four different complexes with critical gene regulatory functions ^12–20^. The canonical functions of two of these complexes are to activate gene transcription whereas the other two most often function in gene repression. The Super Elongation Complex (SEC) including RNA Polymerase II (RNAP) and pTEFb, is directly recruited via interaction of MLLT3/1 with AFF4 (or its homolog AFF1) to promote transcription elongation of paused genes, a rate-limiting step for many mammalian genes that depends on the release of RNAP from the proximal promoter in response to environmental signals ^21^. The second MLLT3/1-recruited complex includes DOT1L, the only H3K79 methyltransferase in animals, whose canonical role is to facilitate active gene transcription ^22^. While the role of DOT1L was originally thought to be via effects on elongation, recent studies show a critical role for DOT1L in transcriptional initiation via its interaction with MLLT3 or MLLT1, with clear reductions in TBP occupancy upon knockdown of DOT1L, MLLT3, or MLLT1 ^23,24^. A third complex is the canonical Polycomb Repressive Complex 1 (PRC1), in which CBX8 mediates the association of MLLT3/1 with other PRC1 components (BMI1, RING1A/B, PH1) ^25–27^. Finally, MLLT3/1 bind to the BCL6 co-repressor protein BCOR, a component of a non-canonical (nc) PRC1 repressor complex which also includes KDM2B which has a non-methyl CpG DNA-binding domain ^28,29^. Current dogma dictates that both PRC1 and ncPRC1 facilitate and reinforce chromatin silencing and compaction. However, more recently context-specific diversity of polycomb functions, particularly with the ncPRC1 relevant to transcriptionally active sites, is starting to be appreciated (reviewed in ^30^).

Loss of H3K9ac via knockdown of GCN5 and PCAF was shown to induce changes in transcriptional elongation for a significant set of genes ^31^. This was linked to loss of SEC binding to the relevant genes. A separate study showed reduced genomic occupancy of MLLT3 in GCN5/PCAF null cells with reduced H3K9ac, again linking MLLT3 and the SEC to H3K9ac binding ^7^. Biochemical measurements have shown a preference of the MLLT3 YEATS domain to bind to H3K9 ac/cr ^7,8,32^. All of this data links the H3K9ac binding of the MLLT3 YEATS domain to its ability to mediate its functional effects on transcription.

7SK is a snRNA that has been shown to be a negative regulator of transcriptional elongation (Nguyen *et al.* 2001, Yang *et al.* 2001). 7SK exists as a ribonucleoprotein (RNP) complex in cells which includes the proteins MePCE, LARP7, HEXIM1/2, and P-TEFb (CDK9 and Cyclin T1/T2) ^33^. P-TEFb kinase activity is required for the phosphorylation of negative elongation factors (NELF and DSIF) as well as Ser2 on RNA Pol II to mediate productive transcriptional elongation. The 7SK complex functions to inhibit transcriptional elongation by inhibiting P-TEFb kinase activity. Both transcription factors as well as coactivators including NF-κB, c-Myc, CIITA, p53, the bromodomain protein 4 (Brd4), the super elongation complexes (SECs), and the human immunodeficiency virus (HIV) transactivator Tat can directly bind to the 7SK RNP complex and target P-TEFb to paused RNAPII ^34,35^.

Recently, it has been demonstrated that a number of epigenetic reader domains also bind nucleic acids ^36^, albeit typically with weak affinity and limited, if any, specificity. Herein we describe our findings that the YEATS domain of MLLT3 is not only a reader of histone acetylation and crotonylation but also binds to ncRNA. Using PAR-CLIP, we identified a number of targets of RNA binding by MLLT3 but among these was the snRNA 7SK. The binding to 7SK is clearly of functional significance considering that MLLT3 is a component of the SEC that enhances transcriptional elongation, as mentioned above. Using biochemical assays, we showed that the MLLT3 YEATS domain binds to SL4 of 7SK with high affinity. Furthermore, we showed that the binding of the YEATS domain of MLLT3 to the SL4 of 7SK competes with the binding of a LARP7 construct containing the RRM and La module domains, suggesting a potential mechanism that could contribute to the release of P-TEFb. Based on mapping of the RNA interaction surface on the YEATS domain using NMR, we developed a mutant form of the YEATS domain which binds 7SK with 10-fold lower affinity but with no effect on histone peptide binding to probe the functional role of the YEATS domain – 7SK interaction. Comparison of the effects of wt and mutant forms of MLLT3 on the differentiation of mouse lineage depleted BM stem/progenitor cells shows a significant skewing of the myeloid lineage upon loss of RNA binding. This mutant form of the protein was used to show by RNA-Seq that loss of RNA binding results in substantial changes in gene expression and by PRO-Seq that loss of RNA binding blocks transcriptional initiation rather than transcriptional elongation. This data establishes the MLLT3 YEATS domain as a dual reader of H3K9ac and the ncRNA 7SK with a critical role in regulation of transcription via effects on both initiation and elongation.

## Results

### MLLT3 YEATS domain binds to RNA in a sequence-dependent manner

We noted in our purification of the MLLT3 YEATS domain that it co-purified with nucleic acid bound. Using a colorimetric assay ^37^, we confirmed that it co-purified with RNA. Based on this, we sought to determine if the MLLT3 YEATS domain binds RNA in a sequence-specific manner. Because one face of the YEATS domain resembles the RNA binding interface of RRM domains, we employed an NMR based approach that has been utilized to define the RNA binding specificity of RRM domains that utilizes chemical shift changes in the ^15^N-^1^H HSQC spectrum of the protein upon addition of a set of short RNA oligomers with systematic variation of the bases ^38^. As shown in Supplementary Figure 1, the data yielded a consensus sequence of uuAUCU or uuAUAU. The third position was the only position at which there were two bases that were equally prevalent. To probe one additional position, we ordered uuAUCUA, uuAUCUU, uuAUCUC, and uuACUCG, each conjugated to a 6-[FAM] fluorophore on the 5’ uracil to facilitate binding measurement by fluorescence polarization. The uuAUCUU oligo showed the highest binding affinity with a K_D_ of 48 nM. This data establishes the MLLT3 YEATS domain as an RNA binding protein.

### MLLT3 binds the ncRNA 7SK

To identify the cellular RNA targets of MLLT3, we employed Fluorescent Photoactivatable Ribonucleoside-Enhanced Crosslinking and Immunoprecipitation (fPAR-CLIP) methodology as previously described ^39^. We selected the HEL cell line for these studies due to its high endogenous MLLT3 expression levels, which allows for 3XFLAG-tagged MLLT3 overexpression without adverse effects on cell growth or viability. Additionally, HEL cells are derived from a patient with erythroleukemia and therefore represent an appropriate hematopoietic model system relevant to MLLT3’s known biological functions.

Following fPAR-CLIP crosslinking and sequencing, nuclear binding peaks were ranked in descending order according to crosslinked read density. This analysis revealed that the small nuclear RNA 7SK contained one of the most robust and highly enriched MLLT3 binding sites identified in our dataset. Detailed examination of Integrative Genomics Viewer (IGV) coverage plots demonstrated a pronounced binding preference for the 3’ region of 7SK, specifically localizing to the stem-loop 4 (SL4) structural domain (Figure 1A-C). This targeted binding pattern suggests that MLLT3 recognizes specific structural features within the 7SK ribonucleoprotein complex rather than exhibiting non-specific RNA association. The known role of 7SK in regulation of transcriptional elongation strongly suggested a functional role for binding of the MLLT3 YEATS domain, thus we pursued further evaluation of the role of RNA binding in MLLT3 YEATS domain function.

**Figure 1.**
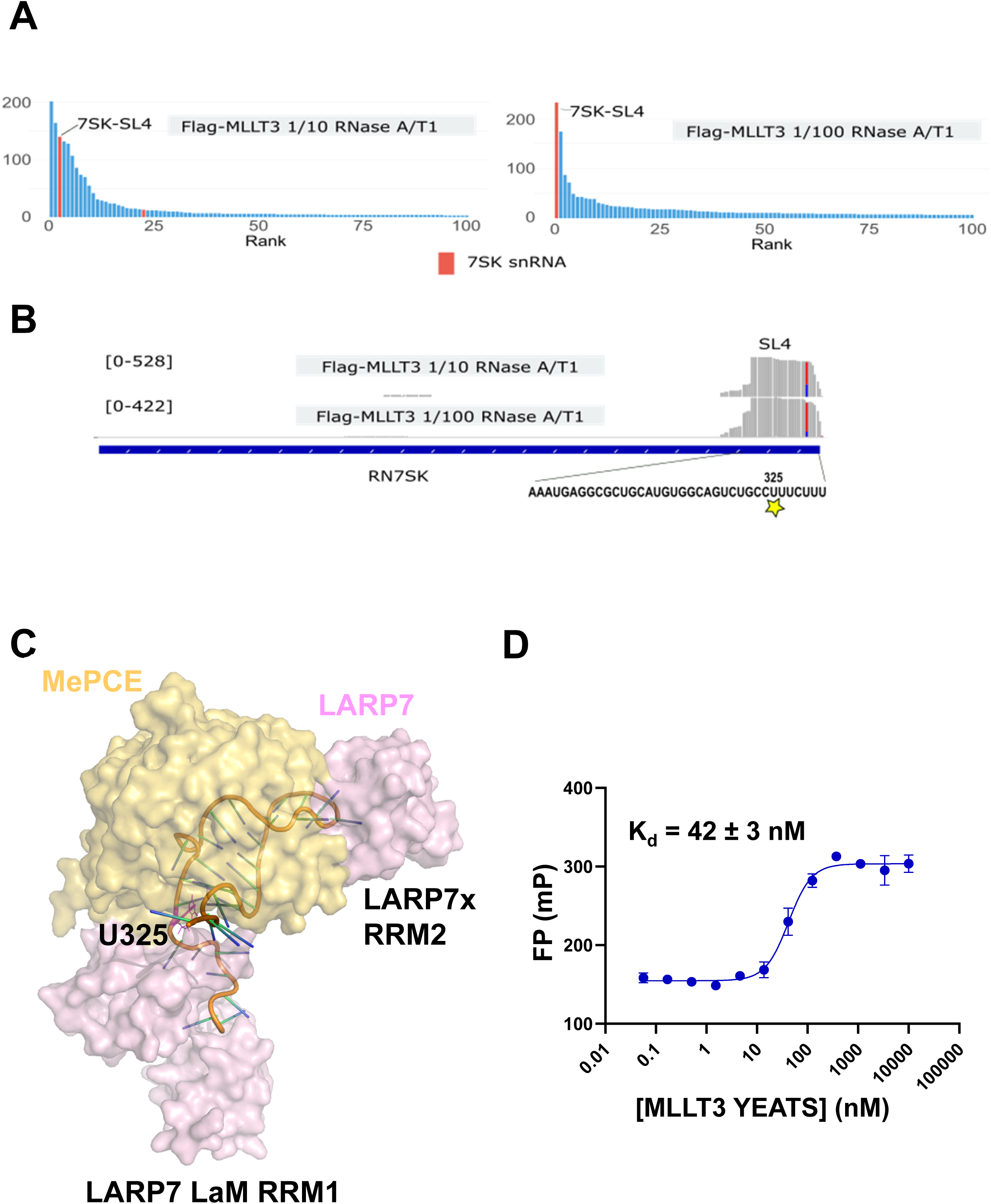
The MLLT3 YEATS domain binds SL4 of 7SK. **A.** Binding sites identified by fPAR-CLIP were ranked according to crosslinked read count for two biological replicates using 1:10 and 1:100 dilutions of RNase A/T1 mix. **B.** Integrative Genomics Viewer (IGV) visualization of aligned sequencing reads derived from fPAR-CLIP experiments. **C.** Surface representation of MePCE and LARP7 bound to stem loop 4 of 7SK (PDB code: 7SLP). RNA is displayed as a cartoon representation with U325, site of covalent attachment to MLLT3 in the PAR-CLIP experiment indicated. **D.** Fluorescence polarization (FP) based assay titrating MLLT3 YEATS domain into Yakima yellow-labeled 7SK SL4. Results shown are from 3 replicates.

The PAR-CLIP approach yields a uracil to cytosine conversion at the sites of RNA cross-linking to protein. Analysis of these sites for cross-linking of MLLT3 to 7SK identified U325, located in the stem loop 4 (SL4) of 7SK, as a site of cross-linking suggesting a binding interaction with SL4 (Figure 1A-C). To biochemically assess the binding of the MLLT3 YEATS domain to 7SK SL4, we used a Yakima yellow labeled 7SK SL4 RNA to measure binding using fluorescence polarization. Direct binding of the MLLT3 YEATS domain yielded a Kd of 42 nM (Figure 1D), indicative of a high affinity interaction. To verify that the fluorescent label does not contribute to the binding, we used a competition experiment competing Yakima yellow 7SK SL4 off the YEATS domain with unlabeled 7SK SL4 (Supplementary Figure 2), showing only a modest inhibitory effect of the fluorophore (K_I_ = 21 nM). For comparison, we have also measured the binding of the homologous ENL YEATS domain to 7SK SL4 and determined a K_d_ of 150 nM (Supplementary Figure 2) showing ENL also can bind 7SK SL4 albeit with lower affinity.

### 7SK SL4 binding site on the MLLT3 YEATS domain

To map the 7SK SL4 binding interface on the YEATS domain, we employed NMR chemical shift perturbation measurements. The backbone NMR resonances of the YEATS domain previously reported ^40^ were confirmed using an HNCACB experiment. ^15^N-^1^H HSQC spectra were recorded for the YEATS domain alone and in the presence of 7SK SL4 and differences in chemical shifts calculated as previously described ^41^ (Figure 2A). These chemical shift changes were mapped onto the structure of the MLLT3 YEATS domain (Figure 2B), showing the RNA binds to a region of the YEATS domain distinct from, but in proximity to, the peptide binding site. Using the sites of NMR chemical shift changes and the identity of the base that is cross-linked to the YEATS domain from our PAR-CLIP data, we docked 7SK SL4 (PDB ID: 7SLP) on the MLLT3 YEATS domain (PDB ID: 5HJB) structure using the HADDOCK webserver. The best model is shown in Figure 2C. This model appears to be self-consistent based on the sites of observed chemical shift changes being in contact with the RNA. To further verify the model and provide a useful reagent to study the biological role of RNA binding to the YEATS domain, we introduced two charge reversal mutations for residues at the RNA binding interface (K63E, K67E). These mutations reduced binding of the YEATS domain to 7SK SL4 16-fold while having no impact on H3K9ac binding (Figure 2D, Supplementary Figure 4). Furthermore, the introduction of these two point mutations did not significantly alter the structure of the YEATS domain, as assessed by comparison of their ^15^N-^1^H HSQC spectra (Supplementary Figure 4E). This further validates the model for binding and provides a high quality reagent to examine the biological role of RNA binding to the YEATS domain. This double mutant was introduced into full-length MLLT3 constructs for functional testing in cells.

**Figure 2.**
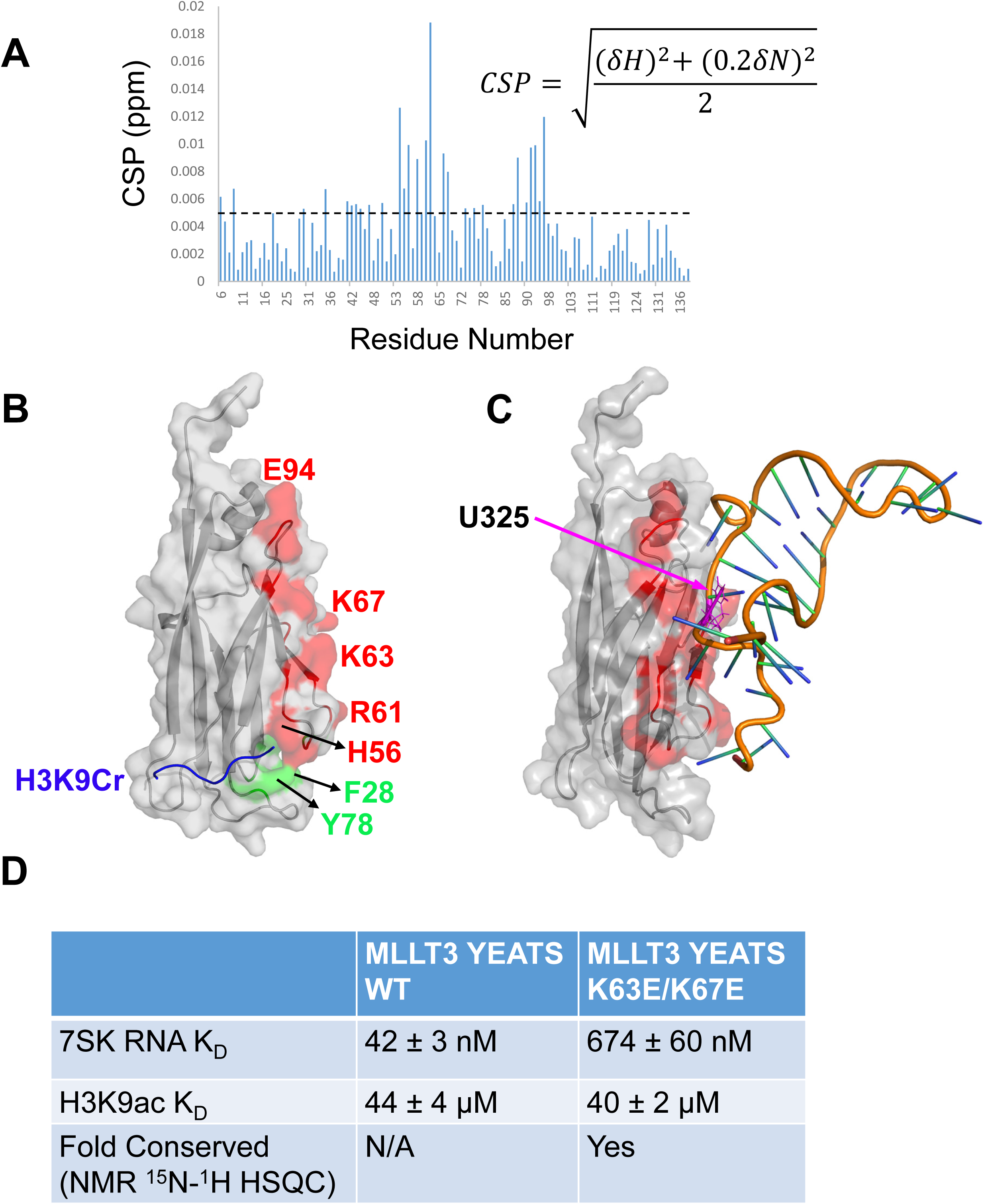
7SK SL4 binds to a site on the MLLT3 YEATS domain distinct from the H3K9ac/cr binding site. **A.** Chemical shift perturbations (CSPs) versus the primary sequence observed in a 2D ^15^N-^1^H HSQC spectrum of the MLLT3 YEATS upon addition of 7SK SL4. **B.** Surface representation of the structure of the MLLT3 YEATS domain (PDB code: 5HJB) with H3K9cr peptide indicated as a coil (blue), sites of mutations that alter H3K9cr binding colored green, and sites of significant chemical shift changes upon addition of 7SK SL4 colored red. **C.** Model of MLLT3 YEATS domain binding to 7SK SL4 determined using HADDOCK. **D.** Table of K_d_ values for wt MLLT3 YEATS domain and K63E/K67E MLLT3 YEATS domain binding to H3K9ac and 7SK SL4 using FP assays. Results shown are for 3 replicates.

### MLLT3 YEATS domain competes with LARP7 for binding to 7SK SL4

Based on a recent structure of 7SK bound to LARP7 and MePCE ^42^, the U325 of 7SK where the PAR-CLIP data shows cross-linking to the YEATS domain is in close proximity to the site where the LARP7 La module and RRM1 binds to 7SK as well as where the Ser530 and Arg531of MePCE makes contacts with 7SK (Figure 3A). Furthermore, our model of binding of the YEATS domain to 7SK SL4 predicts there would be a steric clash between the YEATS domain and LARP7 that suggests the two compete for binding (Figures 1C,3A). To probe whether binding of the MLLT3 YEATS domain can compete with LARP7 for binding, we used a FRET assay with Cerulean-LARP7 (La module + RRM1, aa Met1-Pro208) and Yakima yellow labeled 7SK SL4 RNA (Figure 3B) to first measure the binding constant between the two (K_d_ = 33 ± 1 nM) and then compete with the YEATS domain to determine a K_I_ value (K_I_ = 1846 ± 283 nM). This data establishes that the YEATS domain does compete with LARP7 for binding to the 7SK SL4.

**Figure 3.**
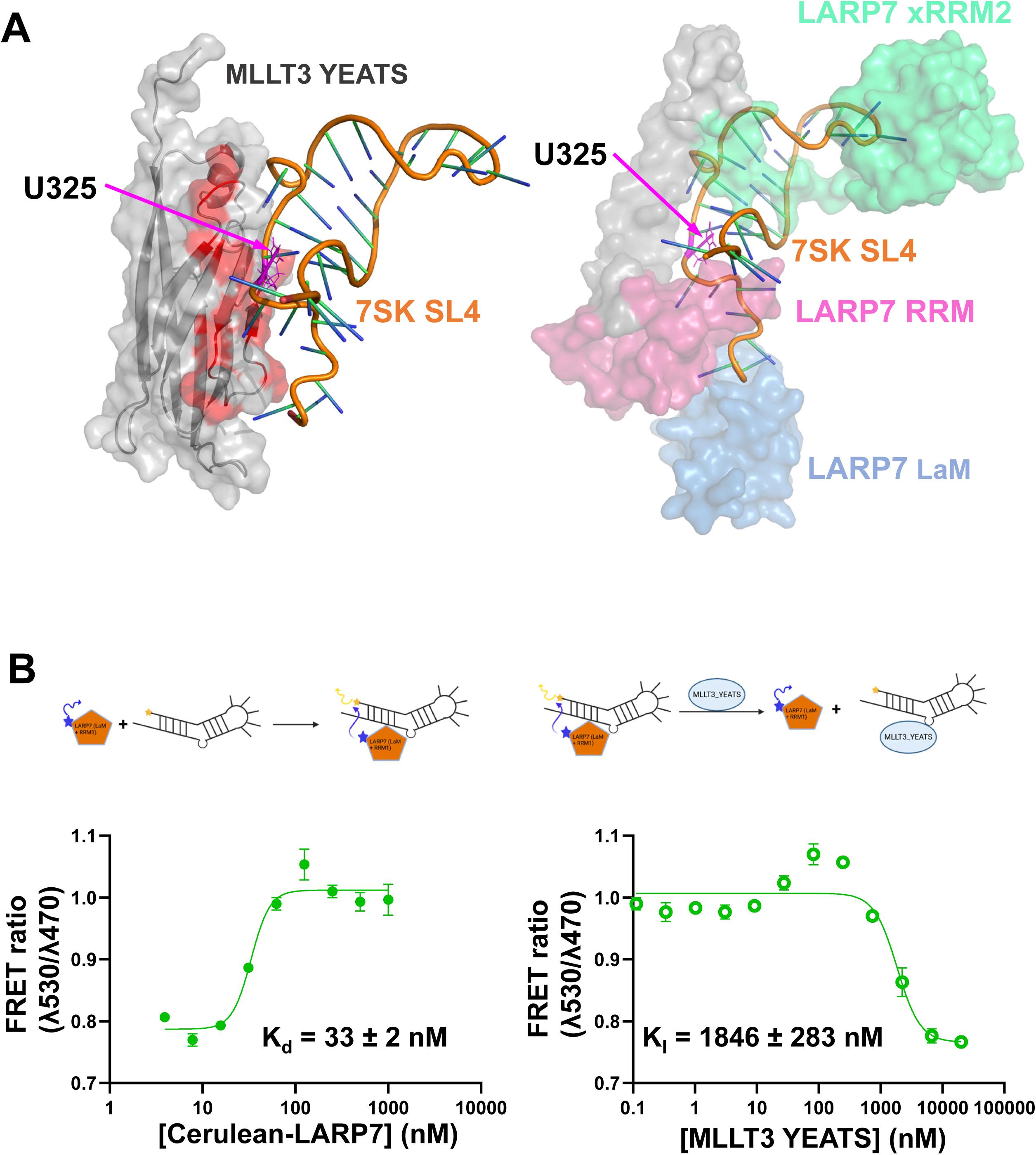
The MLLT3 YEATS domain competes with LARP7 for binding to 7SK. **A.** Left: Model of MLLT3 YEATS domain bound to 7SK SL4. Right: Surface representation of MePCE and LARP7 bound to 7SK SL4 with the U325 site of covalent attachment in the PAR-CLIP indicated, illustrating the overlapping site of binding for the MLLT3 YEATS domain and LARP7. **B.** FRET based competition experiment using Cerulean-LARP7 and Yakima yellow labeled 7SK SL4 and competing with untagged MLLT3 YEATS domain. Left: Determination of the K_d_ for binding of LARP7 to 7SK SL4. Right: Results of a competition experiment where MLLT3 YEATS domain is competing Cerulean-LARP7 off the RNA leading to a loss of the FRET effect.

### Loss of RNA binding to the MLLT3 YEATS domain decreases monocytic differentiation and increases the CLP population

To examine whether the RNA binding function of the MLLT3 YEATS domain is relevant to hematopoiesis, we introduced wildtype (wt) and K63E/K67E double mutant (dm) MLLT3 into lineage-depleted murine BM cells, in conjunction with knockout of the endogenous *Mllt3* gene, to provide a clean genetic background. Comparison of the effects of wt and mutant forms of MLLT3 on the differentiation of mouse lineage depleted BM stem/progenitor cells shows a significant effect on the myeloid lineage upon loss of RNA binding. This was most predominant on the monocytes (LY6Chi, CD11b int, GR1 low/int) population, which was dramatically decreased (Fig 4A (left panels), 4B). This was in contrast to minimal change in granulocyte/macrophage populations (CD11bhi) (Fig. 4A (left panels)), and an increase in common lymphoid progenitor population (CLP) (Fig. 4A (right panels), 4B). Decreased proliferation and expansion was observed in the DM cells as well (Fig. 4C).

**Figure 4.**
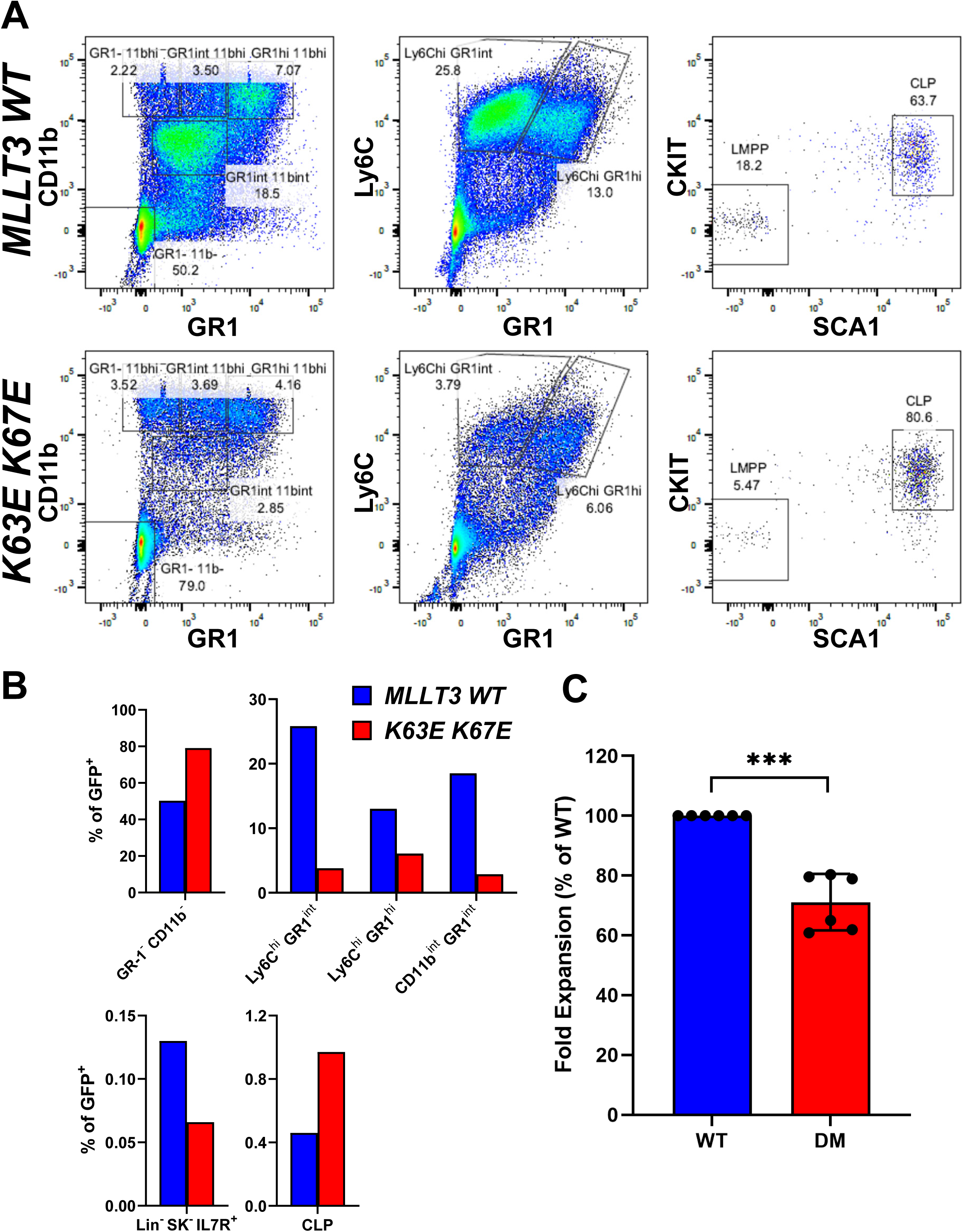
Loss of RNA binding of MLLT3 alters hematopoiesis. **A.** Surface marker staining of cultured *Mllt3* KO bone marrow progenitor cells overexpressing MLLT3 WT or DM indicating a reduction of monocyte and granulocyte populations and an increase in common lymphoid progenitors in the DM sample. Staining was performed on a pooled sample comprised of three biologic replicate samples. **B.** Bar graphs of data shown in **A**. **C.** Relative cell expansion of cells in culture. Results shown are from 6 replicates.

### Loss of RNA binding to the MLLT3 YEATS domain abrogates the transforming properties of an NES-MLLT3 construct

The functionality of MLLT3 and MLLT1 can be evaluated by their contribution to leukemic transformation of hematopoietic progenitors. The CALM-AF10 fusion gene transforms hematopoietic progenitors through the functions mediated by the NES and the OM-LG domain which binds to DOT1L and AF9/ENL proteins. Its leukemic transformation can be mimicked by an artificial fusion gene construct in which the NES of CALM is fused to the entire coding sequence of MLLT3 or MLLT1^43^. Leukemic transformation by leukemia oncogenes can be evaluated using an *ex vivo* myeloid progenitor transformation assay ^44,45^(Figure 5A). Cells transduced with a functional oncogene typically exhibit upregulated expression of *Hoxa9* in first-round colonies and vigorous colony-forming activity in the third and fourth rounds of replating. The NES-MLLT3 construct produced colonies at the third and fourth rounds of plating and exhibited high level *Hoxa9* expression as previously reported. Introduction of a Y78A mutation in the YEATS domain of MLLT3 resulted in loss of transformation, indicating that the recognition of acylated histone H3K9/18/27 is required for transformation. Next, we introduced the K63E/K67E double mutations in the NES-MLLT3 construct (KKEE), which failed to transform hematopoietic progenitors. This indicates that the RNA binding function of the MLLT3 YEATS domain is essential for transformation.

**Figure 5.**
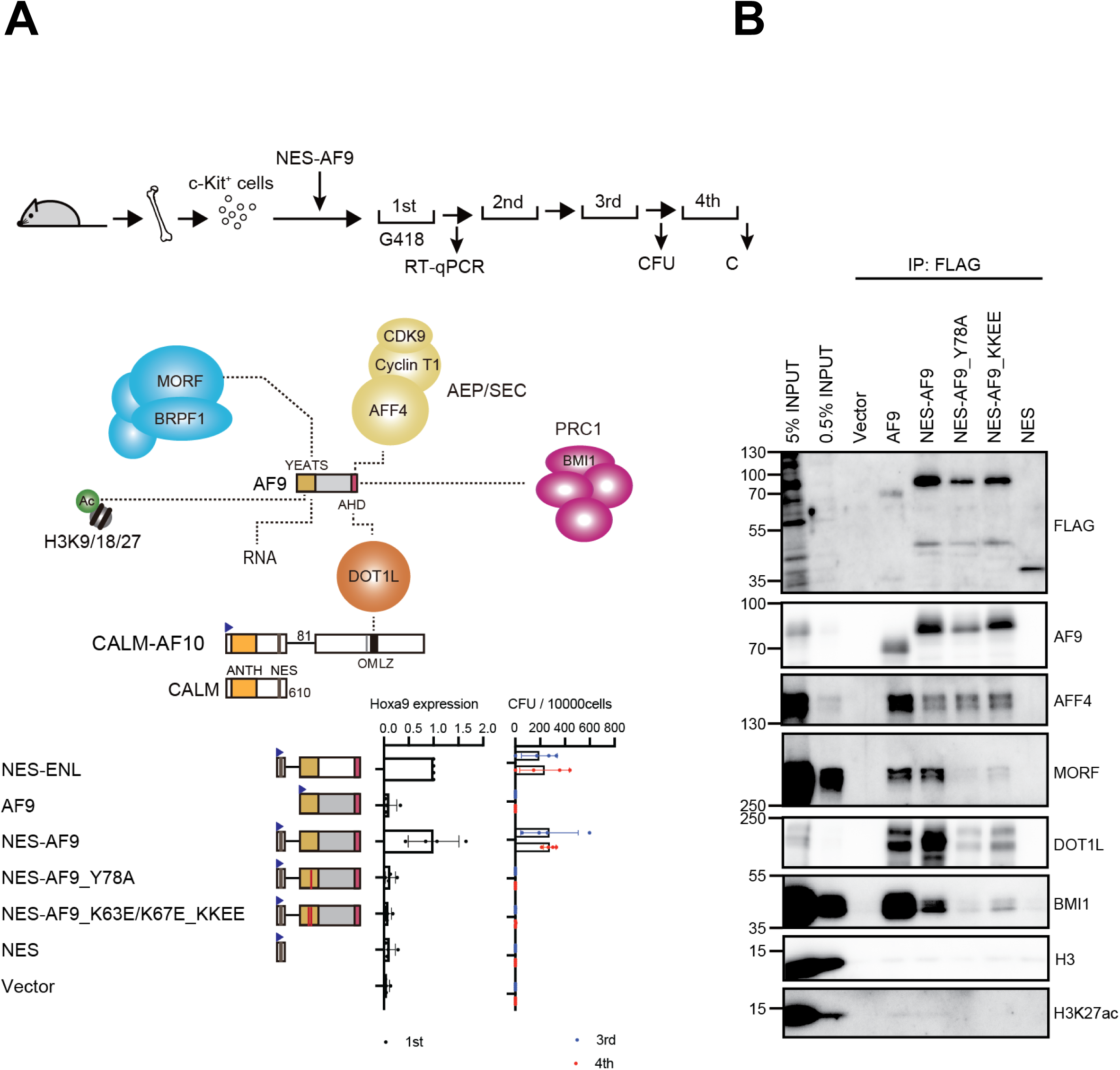
Loss of RNA binding abrogates the transformation capability of NES-MLLT3. **A.** Oncogenic activity of NES-AF9 fusion. Various NES-AF9 fusions were tested for their oncogenic activities using myeloid progenitor transformation assay^45^, outlined on top. Colony forming units (CFU/10,000cells) from three biological replicates are shown with error bars (SD). Hoxa9 expression (normalized to Gapdh) in the first-round passage is analyzed by RT-qPCR (three biological replicates with SD). **B.** Protein interaction of NES-AF9 fusions on chromatin. HEK293T cells were transiently transfected with various FLAG-tagged NES-AF9 fusions carrying K63E/K67E mutations that abrogate RNA interaction and analyzed by the fanChIP method^46^ using anti-FLAG antibody (Y78A mutation, which abrogates YEATS-acyl-lysine interaction, was included for comparison). The K63E/K67E mutations reduced the interaction with MORF, DOT1L and BMI, whereas AFF4 interaction was unaffected.

We also examined protein-protein interactions on chromatin by the NES-MLLT3 construct using the fractionation-assisted chromatin immunoprecipitation (fanChIP) method^46^ (Figure 5B). NES-AF9 associated with the co-factors previously reported including AFF4, DOT1L, BMI1, and MORF. The Y78A mutant was detected in a much smaller amount compared to other NES-MLLT3 constructs and showed a reduced binding of its co-factors except AFF4. The KKEE mutant was detected in a comparable amount to its wildtype counterpart, but lost most of its MORF binding, and showed markedly reduced binding to DOT1L and BMI1. These results suggest that RNA binding by the YEATS domain may be necessary for the efficient assembly of co-factors on chromatin.

### Loss of RNA binding by the MLLT3 YEATS domain alters target gene expression

To understand the mechanism of altered hematopoiesis with loss of MLLT3 RNA binding, we analyzed expression in MLLT3 wt and dm expressing mouse lin-BM cells following in vitro culture (as in Figure 4) using RNA-seq. A total of 1409 differentially expressed genes were identified (FC > 1.5, p-adj <0.05). A volcano plot (Fig. 6A) shows that the most substantive changes are observed for genes that are reduced in expression. Decreased DNA replication and cell cycle KEGG pathways corroborate the observed decreased proliferation and cell expansion (Figure 4C). GSEA showed significant enrichment of gene sets relevant to DNA replication, cell cycle, and multiple hematopoietic lineages (Supplemental Fig. 6A,B) which aligns with RNAseq KEGG pathway analysis (Fig. 6C). Notably, numerous members of the *Hoxa* cluster (*Hoxa4*,*5*,*6*,*7*,*9*,*10*) are substantially reduced in expression, likely contributing to the observed decreased cell expansion (Fig. 4C,6B). In line with the myeloid lineage phenotype observed, *Ly6c2*, *Ly6g*, *Cebpe*, *Gfi1*, *Epx*, *Elane*, among others, are significantly decreased. In contrast, multiple GSEA pathways and genes relevant to lymphoid development were increased (Fig. 6B, Supp Fig. 5,6B) including *IL7r*, *Cd74*, *Cd180*, *Ly9* and *Cd22*. Together, these data suggest that the MLLT3 YEATS domain RNA binding activity is critical to maintain appropriate gene expression required for normal hematopoietic differentiation programs.

**Figure 6.**
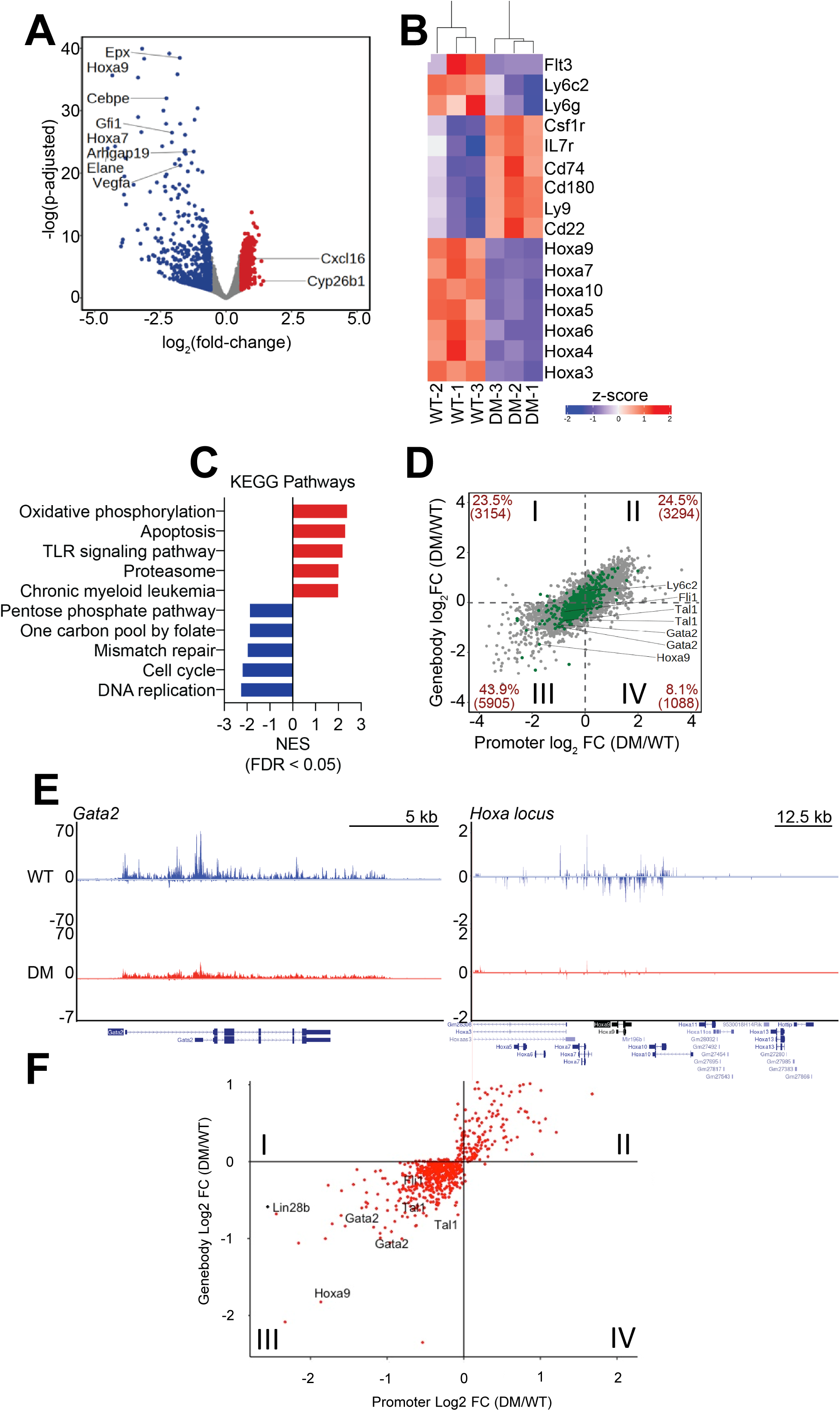
Effects of loss of RNA binding of the MLLT3 YEATS domain on gene expression. **A.** RNAseq comparison of gene expression from cultured *Mllt3* KO bone marrow cells overexpressing WT and K63E/K67E MLLT3. Volcano plot with selected genes indicated. **B.** Heatmap of changes in gene expression for selected genes between wt and K63E/K67E MLLT3 **C.** KEGG pathways identified as significantly altered for wt versus K63E/K67E MLLT3. **D.** Traveling matrix representation of the PROseq data for wt and K63E/K67E MLLT3. MLLT3 target genes from ChIPseq ^2^ are indicated in green. **E.** Selected tracks from the PROseq data for *Gata2* and the *Hoxa* cluster. **F.** MLLT3 target genes plotted on the PROseq traveling matrix colored according to whether a 7SK binding site was identified in or near the gene (red) or no 7SK binding site was found in or near the gene (black). All but one gene, *Lin28b*, had a 7SK binding site in or near the gene.

Based on the established roles of the SEC and 7SK in transcriptional elongation, we expected an effect on elongation so we also carried out precision run-on and sequencing (PRO-seq) using the same wildtype and mutant cells employed for the hematopoietic differentiation and RNA-seq experiments. Alterations to transcriptional elongation were assessed using the traveling matrix analysis ^47^ as shown in Figure 6D. Comparing dm to wt, we see altered genes populating quadrants II and III, with the largest changes occurring in genes in quadrant III. Quadrant III corresponds to loss of RNA PolII both at the promoter and in the gene body. This is consistent with a loss of transcriptional initiation rather than a defect in transcriptional elongation. Comparison of TBP occupancy ^48^ for the class II and III genes that are direct targets ^2^ shows a strong enrichment for the class III genes relative to the class II genes (Supp Fig. 7), consistent with productive initiation. Fig. 6D also highlights genes identified as targets of MLLT3 using ChIP-Seq ^2^ on the traveling matrix plot. We compared 7SK occupancy from ChIRP-Seq data ^49^ to the MLLT3 ChIP-Seq targets on the traveling matrix (Fig. 6F). This analysis showed all but one (*Lin28b*) of the MLLT3 target genes has a 7SK binding site in or near the gene.

**Figure 7.**
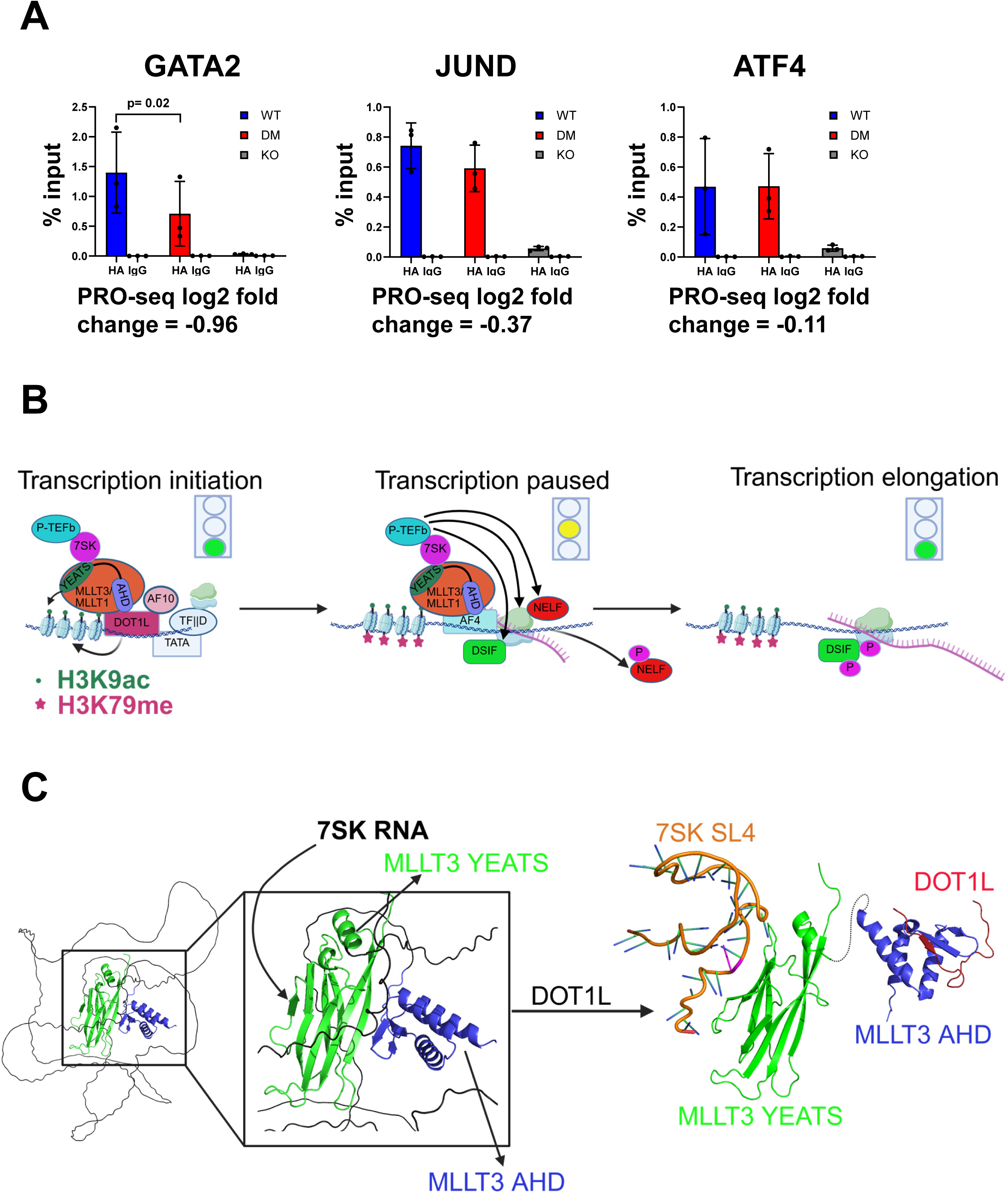
Model of MLLT3 function in transcriptional initiation and elongation. **A.** Results of ChIP qPCR for HA-tagged wt and K63E/K67E MLLT3 on 3 genes with descending effects in the PROseq data. **B.** Proposed model of the mechanism of MLLT3 regulation of transcriptional initiation and elongation. **C.** Proposed model of structural changes in MLLT3 upon 7SK binding.

ChIP-qPCR for wt and K63EK67E MLLT3 on 3 of these quadrant III genes that are known binding sites for MLLT3 (*GATA2*, *JUND*, *ATF4*) shows that *GATA2* occupancy is significantly reduced upon loss of RNA binding whereas *JUND* and *ATF4* were not (Figure 7A). This correlates well with the magnitude of the changes in the PRO-seq data for these three genes where *GATA2* shows a large change whereas *JUND* and *ATF4* show only modest changes.

### Loss of RNA binding reduces MLLT3 occupancy at target genes

HEL cells with MLLT3 KO followed by lentiviral transduction to express HA-tagged-WT or –DM MLLT3 were used for ChIP-q-PCR with anti-HA or IgG antibodies. Three genes that are validated targets of MLLT3 binding and which showed decreasing magnitudes of effects in the PRO-seq were analyzed (GATA2 > JUND > ATF4). Consistent with the effects seen in the PRO-seq data, occupancy of MLLT3 at GATA2 showed a statistically significant loss of occupancy with loss of RNA binding, MLLT3 at JUND showed a trend toward reduced occupancy, and MLLT3 at ATF4 did not show a change in occupancy.

## Discussion

Previous studies showed that the AF9 (MLLT3) YEATS domain binds to acetylated and crotonylated histone peptides ^7,8^ as well as to acetylated MOZ ^50^. Our results show that MLLT3 is both an epigenetic reader as well as a reader of RNA, placing it in a unique class of proteins that are able to bind to both ^36^. Incorporating both functions into a single protein makes it possible for these two signaling pathways to communicate. The dual functionality of the YEATS domain will bind sites in the genome where H3K9ac/cr (or MOZ K1007ac) and 7SK are present and therefore serves as a selection filter for MLLT3 activity. Binding to 7SK in particular, with its well-documented role in transcriptional elongation, makes functional sense for a protein that is a component of the super elongation complex (SEC). Our data show a strong overlap between MLLT3 and 7SK binding sites in the genome, consistent with a functional interaction. The ability of the MLLT3 YEATS domain to compete with LARP7 for binding to 7SK suggests that MLLT3 could modulate 7SK function. Precedent for a regulator of hematopoiesis binding to 7SK comes from the recent finding that IGF2BP3 binds 7SK to regulate megakaryocyte development ^51^. Indeed, the binding site for IGF2BP3 also involves SL4 on 7SK. The interface on the MLLT3 YEATS domain that we identified as the site for RNA binding was recently suggested to be a TFIID interaction surface based on a deletion analysis ^52^. While possible, it seems unlikely both TFIID and 7SK could bind this site. Furthermore, the deletion utilized (one entire β-strand in a β-sandwich structure) would almost certainly result in complete disruption of the 3D structure of the YEATS domain so trying to assign functional effects to this region of the protein based on this deletion may not be informative. Both MLLT3 and its close homolog MLLT1 have been shown to form condensates ^52,53^. Condensates often rely on RNA components for condensate formation, so a possible explanation of the observed phenotype is the loss of 7SK binding reduces the condensate behavior of MLLT3 and this loss prevents TFIID binding. A Poly-Ser region in the intrinsically disordered region (IDR) of MLLT3 was shown to be essential for self-association ^52^. Deletion of this Poly-Ser region led to loss of self-association as well as loss of TBP binding to target genes, consistent with this proposed explanation. Future efforts to understand whether the RNA binding function of MLLT3 influences condensate behavior and what the functional effects are will be important.

The identification of point mutations which can disrupt RNA binding but not H3K9ac/cr binding to the YEATS domain provided us with a biological reagent that could specifically probe the importance of RNA binding to the YEATS domain of MLLT3. The loss of RNA binding causes a substantial effect on hematopoiesis, with a skewing of cells away from the myeloid lineage and toward the lymphoid lineage, establishing a clear role for the RNA binding function of MLLT3 in hematopoiesis. The cellular differences were reflected in significant gene expression changes highlighted by KEGG and GSEA pathways that included programs related to replication, cell cycle, and hematopoietic differentiation. NES-MLLT3 fusion constructs are transforming, resulting in enhanced colony formation and *Hoxa9* expression ^43^. Disrupting RNA binding abrogated both colony formation and *Hoxa9* expression, providing an independent functional context where RNA binding is essential. Interestingly, introduction of the mutations that disrupt RNA binding in this context also disrupted DOT1L recruitment. AlphaFold prediction of the structure of full-length MLLT3 (Fig. 7C) shows a structure where the YEATS domain and the C-terminal AHD are bound to one another. The predicted binding mode has a β-strand from the YEATS domain binding to the AHD in a manner such that partner proteins (AFF1 (AF4), DOT1L, BCOR, CBX8) could not bind ^18,20,54^. This would lead to inhibition of partner protein binding, i.e. the protein could be regulated by autoinhibition. Binding of the 7SK RNA could displace the AHD and release it to bind to partners including DOT1L.

Disruption of RNA binding by the YEATS domain of MLLT3 results in substantial changes in gene expression. Changes in specific genes (*Ly6c2*, *Ly6g*, *Cebpe*, *Gfi1*, *Epx*, *Elane* for example) are consistent with the shift we see from the myeloid to the lymphoid lineage in hematopoiesis assays. Because of the role of MLLT3 and the super elongation complex (SEC) more generally in regulating transcriptional elongation, we also carried out a PRO-seq study. This data showed that disruption of RNA binding resulted in loss of RNA PolII at <u>both</u> the promoter and the gene body, consistent with a defect in transcriptional initiation rather than transcriptional elongation. This is consistent with a number of recent reports. Knockdown of MLLT3 was shown to reduce TBP occupancy on MLLT3 target genes ^52^. As noted above, TBP occupancy was enriched for our Class III direct MLLT3 target genes. For the close homolog MLLT1, knockdown of MLLT1 was shown to dramatically reduce TBP occupancy genome-wide ^24^. These authors further showed that DOT1L plays a critical role in transcriptional initiation. As DOT1L is a binding partner for MLLT3 and MLLT1, the data suggest binding of DOT1L to MLLT3 (or MLLT1) mediates productive transcriptional initiation via recruitment of TBP. Other studies identified a Ser rich domain in AFF1 (AF4) that interacts with the 7SK SL1 to load TBP ^55,56^ and showed a functional role for this interaction in MLL fusion driven transcription. Like DOT1L, AFF1 is also a binding partner of MLLT3 and MLLT1, providing another possible pathway for TBP loading. Based on their results, the authors suggested this transcriptional initiation step was rate-limiting for gene activation.

Combining our results with previous studies, we propose the model depicted in Figure 7 for gene activation by MLLT3. Binding of MLLT3 is autoinhibited via interactions between the YEATS domain and the C-terminal ANC1 homology domain (AHD). Binding to sites in the genome with 7SK and H3K9ac/cr results in release of autoinhibition and the AHD being able to bind partner proteins. Binding of DOT1L to the AHD leads to recruitment of TBP (TFIID) to effect productive transcriptional initiation. Subsequently, AFF1 (AF4) is recruited to effect productive transcriptional elongation. We favor this order of events as studies of MLLT1 condensates show that the CDK9 containing MLLT1 condensates on their own exhibit dynamics consistent with a static solid-like state that is unlikely to be favorable for CDK9 activity on RNA PolII, NELF, and DSIF whereas the inclusion of AFF1 (AF4) in the droplet resulted in liquid-like behavior which should be favorable for CDK9 activity on RNA PolII, NELF, and DSIF. Based on these results, introduction of AFF1 as the last step would seem likely.

## Acknowledgements

This work was supported by NIH R01CA233749 awarded to JB, NZ, and MF. AK was supported by a University of Virginia Comprehensive Cancer Center (UVACCC) Trainee Fellowship. We acknowledge support from the University of Virginia Comprehensive Cancer Center (UVACCC) supported Bioinformatics Core, RRID:SCR_012718. This research was supported in part by the Intramural Research Program of the National Institutes of Health (NIH). The contributions of the NIH authors were made as part of their official duties as NIH federal employees, are in compliance with agency policy requirements, and are considered Works of the United States Government. However, the findings and conclusions presented in this paper are those of the authors and do not necessarily reflect the views of the NIH or the U.S. Department of Health and Human Services.

## Data availability

The RNA-seq, PRO-seq, and PAR-CLIP data have been deposited at GEO under superseries GSE303306.

## Supplementary Figure Legends

**Supp Figure 1.**
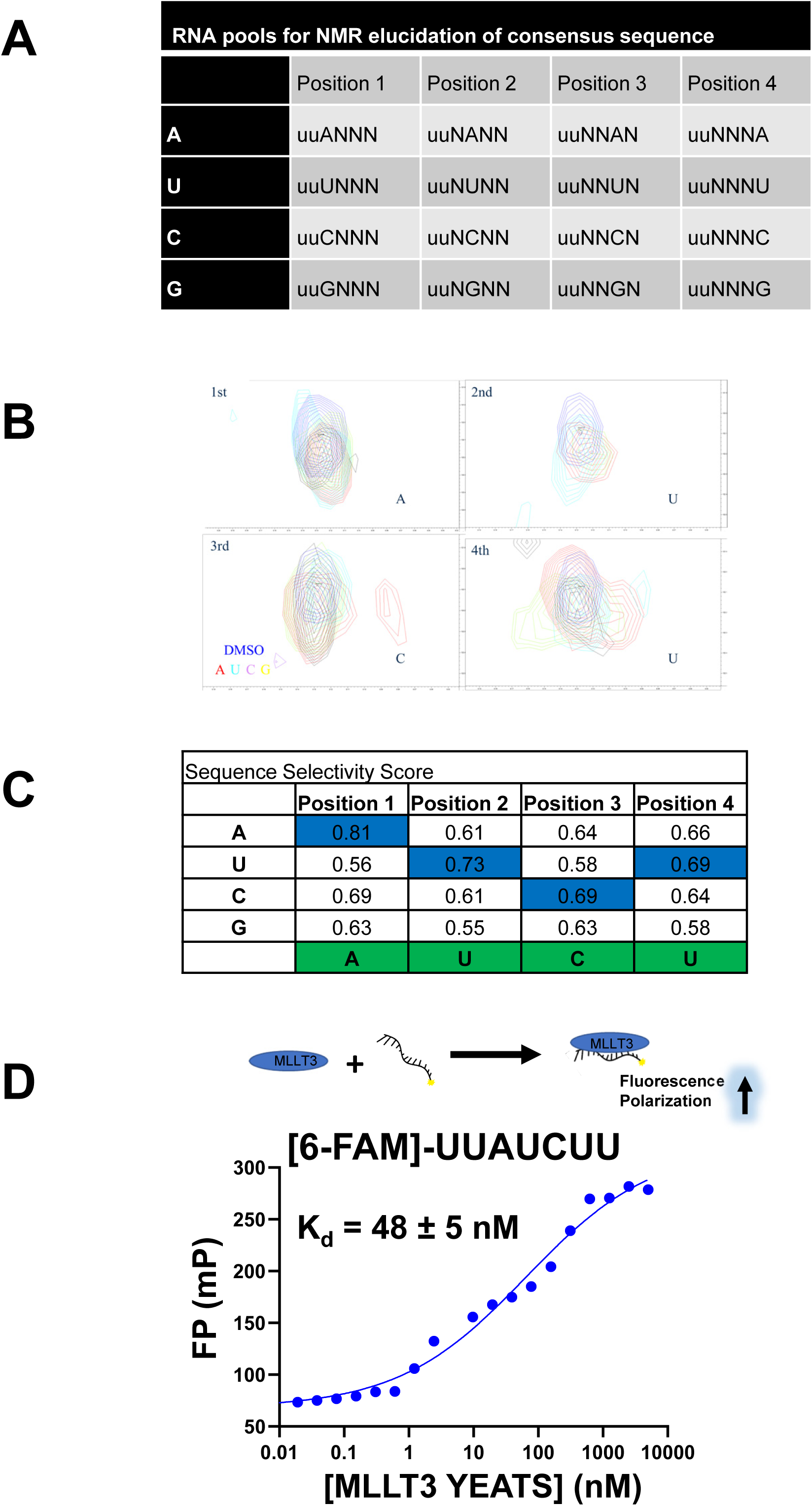
NMR based identification of an RNA binding motif for the MLLT3 YEATS domain. **A**. Table of RNA pools used for the NMR screen. **B.** Chemical shift perturbations observed for selected peaks in the ^15^N-^1^H HSQC of the MLLT3 YEATS domain upon addition of RNA oligos. **C.** Sequence selectivity score for the NMR screen and the RNA motif identified (highlighted in green). **D.** FP assays for binding of the MLLT3 YEATS domain to fluorescein-labeled UUAUCUU.

**Supp Figure 2.**
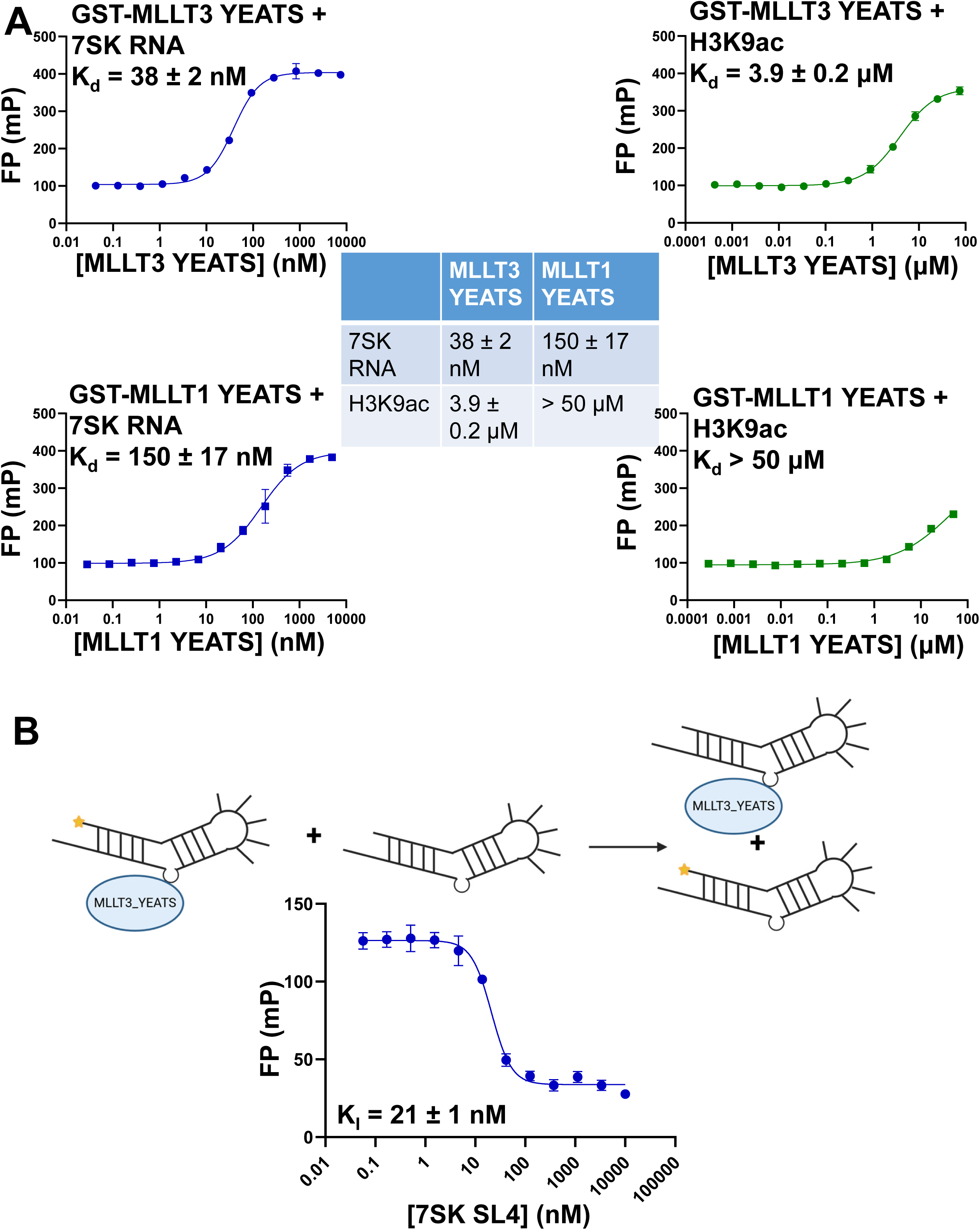
Comparison of MLLT3 and MLLT1 YEATS domain binding to 7SK SL4 and H3K9ac (related to Figure 1) **A.** Top: Results of FP assays for binding of GST-MLLT3 YEATS domain with 7SK SL4 and H3K9ac. Results shown are from 3 replicates. Bottom: Results of FP assays for binding of GST-MLLT1 YEATS domain with 7SK SL4 and H3K9ac. Results shown are from 3 replicates. **B.** Results of competition FP assay with unlabeled 7SK SL4 competing Yakima yellow-labeled 7SK RNA off the MLLT3 YEATS domain. The calculated K_I_ value is shown.

**Supp Figure 3.**
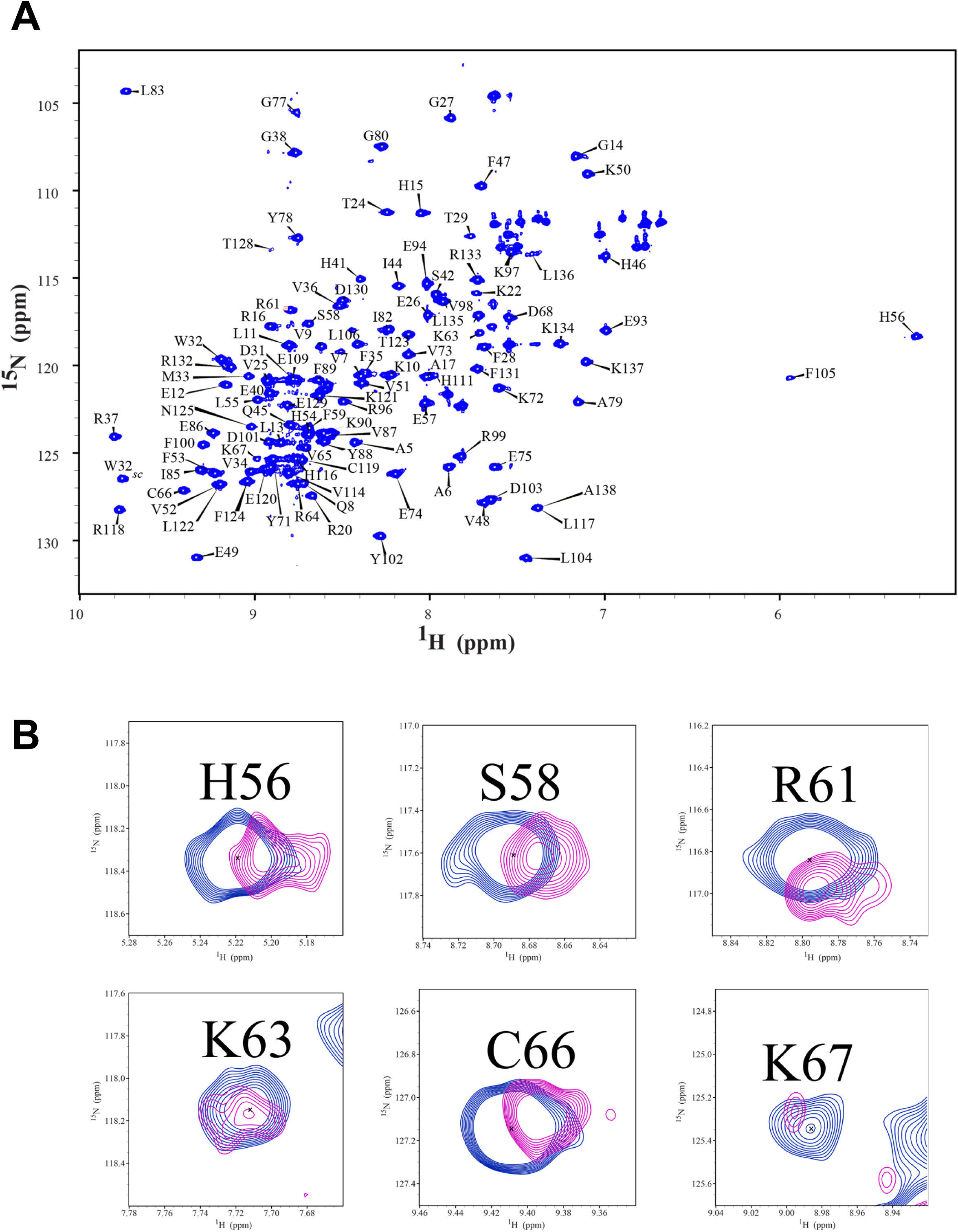
NMR data for wt MLLT3 YEATS domain (related to Figure 2) **A.** ^15^N –^1^H HSQC NMR spectrum of wt MLLT3 YEATS domain. **B.** Selected peaks from the ^15^N-^1^H HSQC spectrum of the MLLT3 YEATS domain showing chemical shift perturbations upon addition of 7SK SL4.

**Supp Figure 4.**
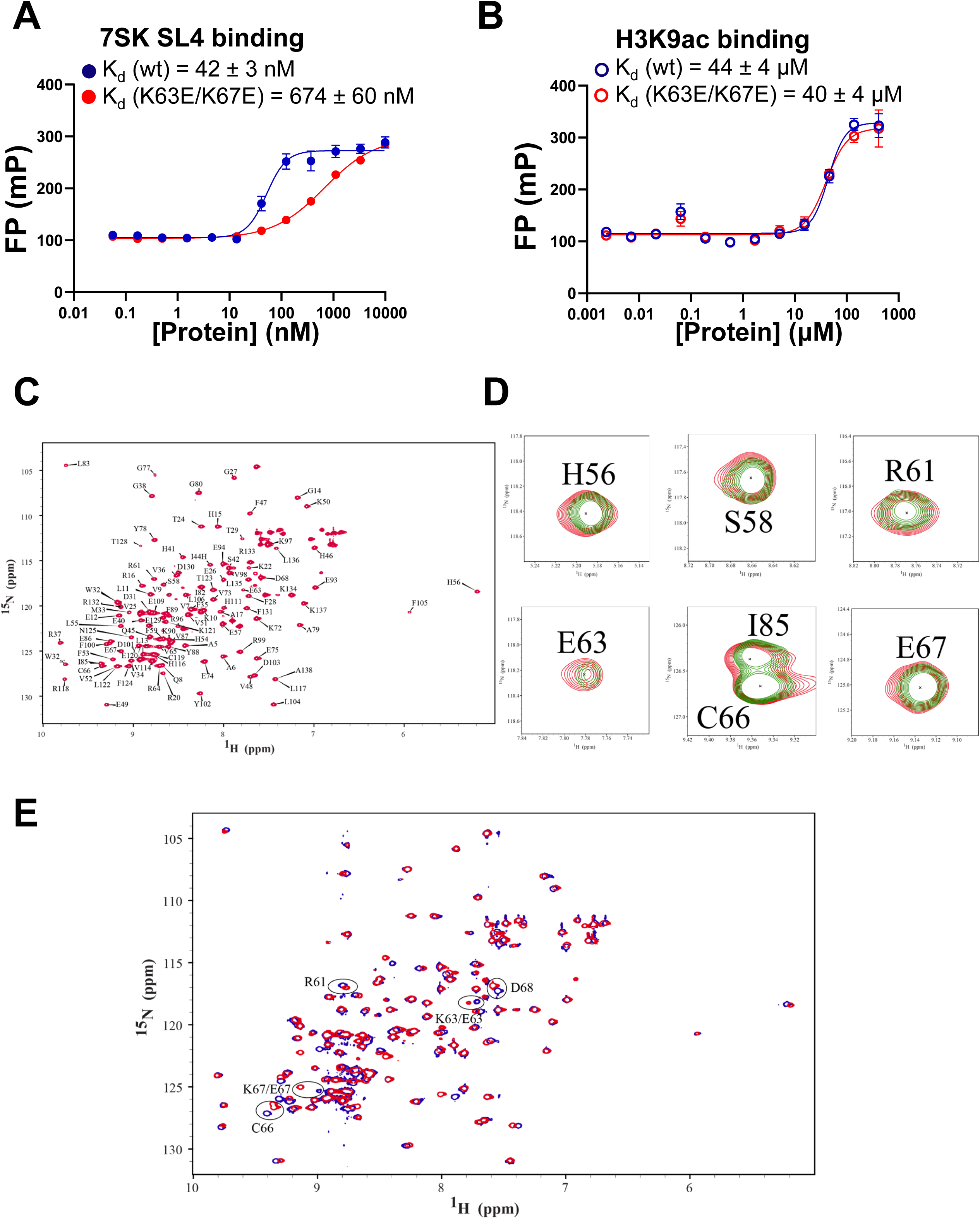
Assay and NMR data for K63E/K67E MLLT3 YEATS domain (related to Figure 2) **A.** FP assay data for binding of K63E/K67E MLLT3 YEATS domain to 7SK SL4. Results shown are from 3 replicates. **B.** FP assay data for binding of K63E/K67E MLLT3 YEATS domain to H3K9ac. Results shown are from 3 replicates. **C.** ^15^N –^1^H HSQC NMR spectrum of K63E/K67E MLLT3 YEATS domain. **D.** Selected peaks from the ^15^N-^1^H HSQC spectrum of the K63E/K67E MLLT3 YEATS domain showing no chemical shift perturbations upon addition of 7SK SL4 (same peaks selected as for Supp Figure 3). **E.** Overlay of ^15^N-^1^H HSQC spectra of wt and K63E/K67E MLLT3 YEATS domains.

**Supp Figure 5.**
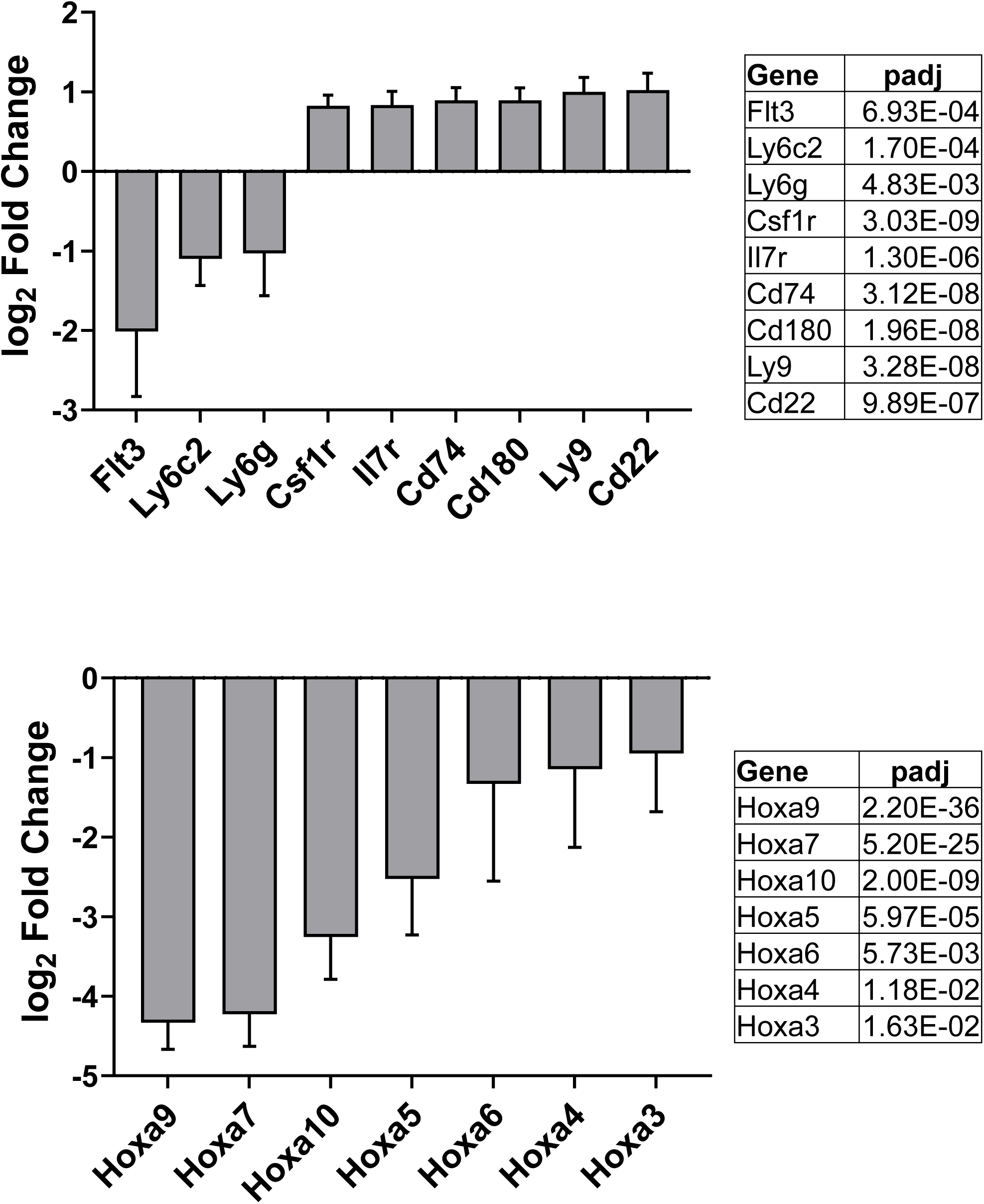
Changes in expression of select genes (related to Figure 6B) Log_2_ fold changes and padj values for selected genes from the RNAseq data.

**Supp Figure 6.**
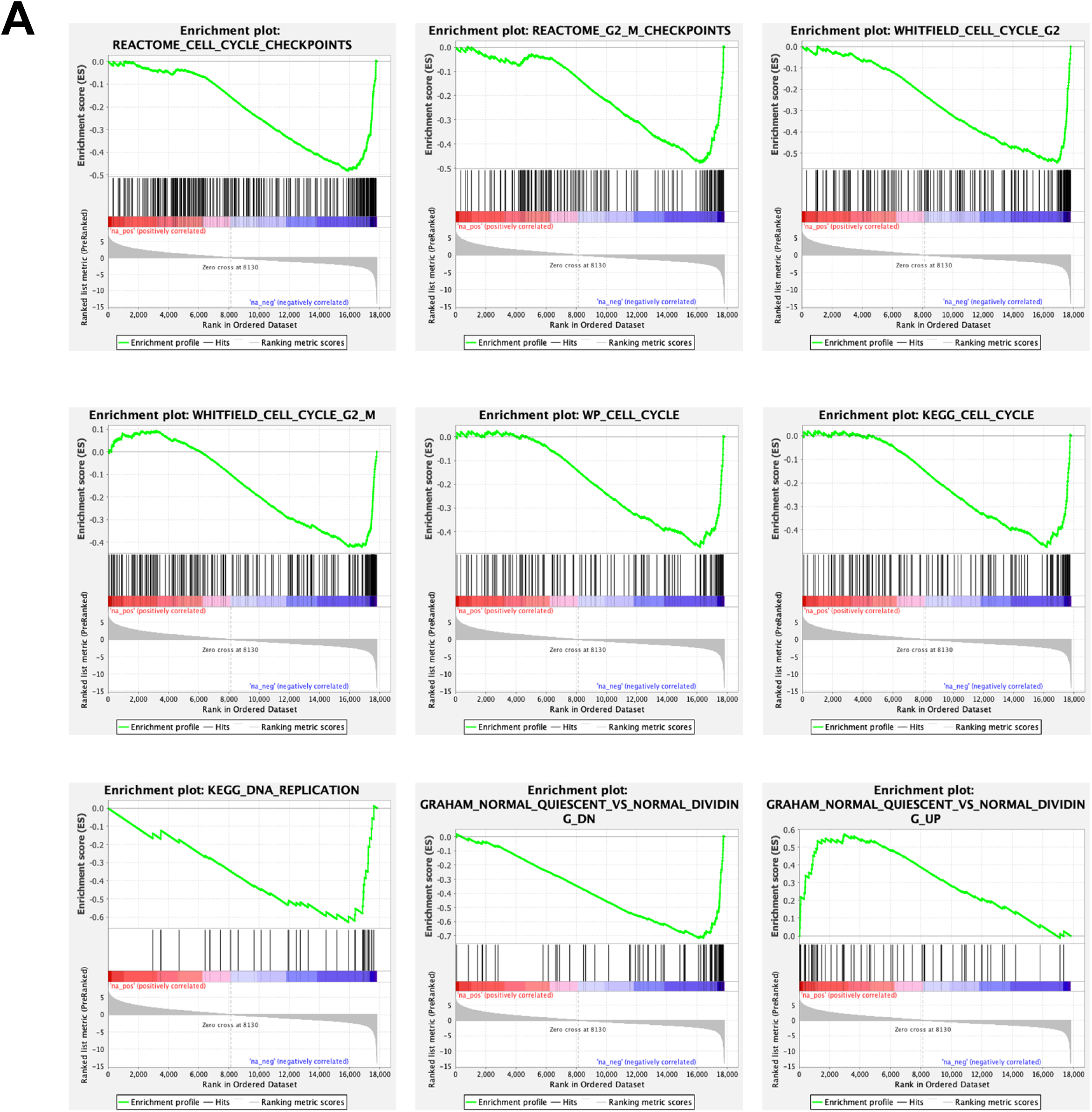
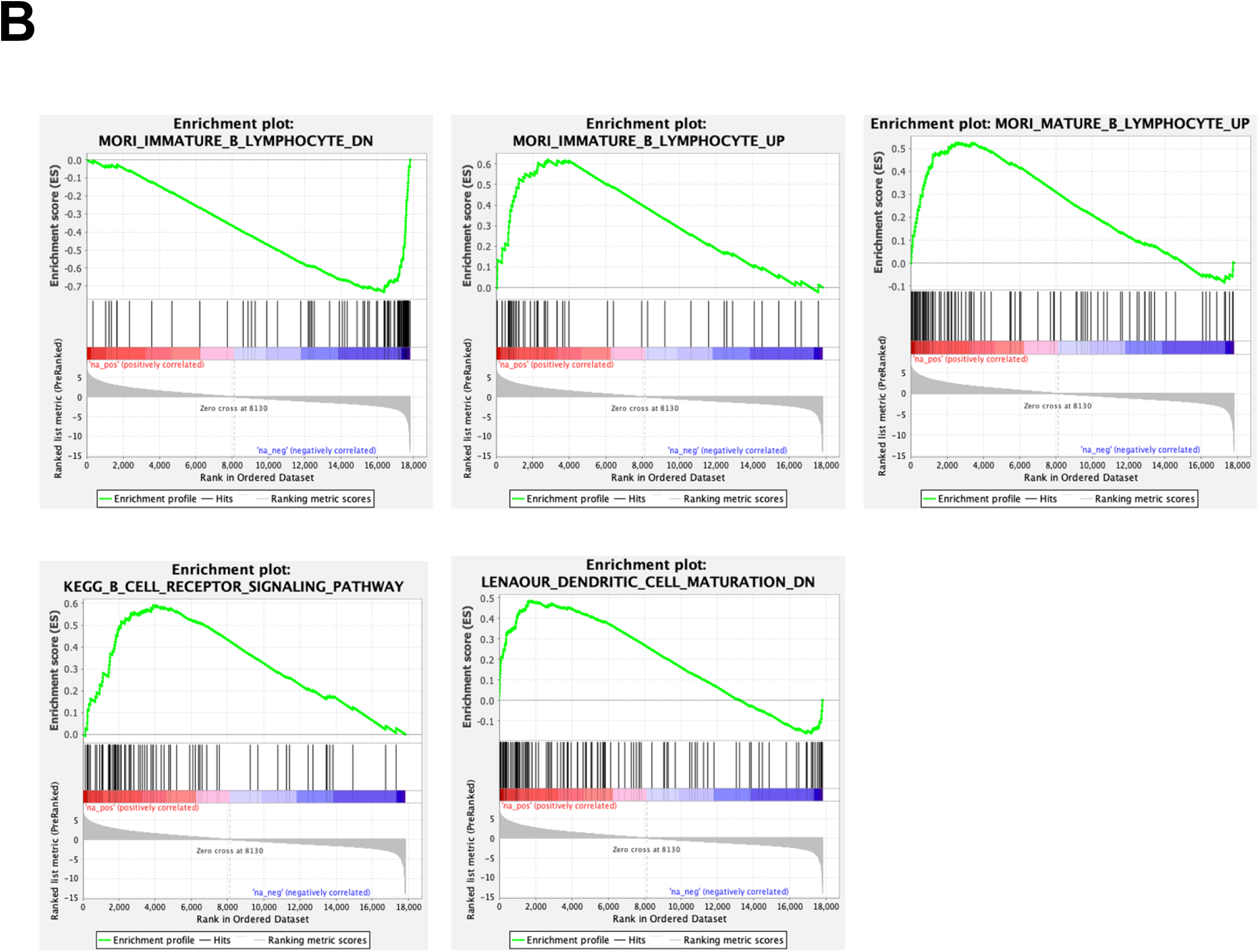
GSEA plots based on the RNAseq data (related to Figure 6) **A.** Gene set enrichment for datasets related to cell cycle, replication, and quiescence. **B.** Gene set enrichment for datasets related to hematopoiesis.

**Supp Figure 7.**
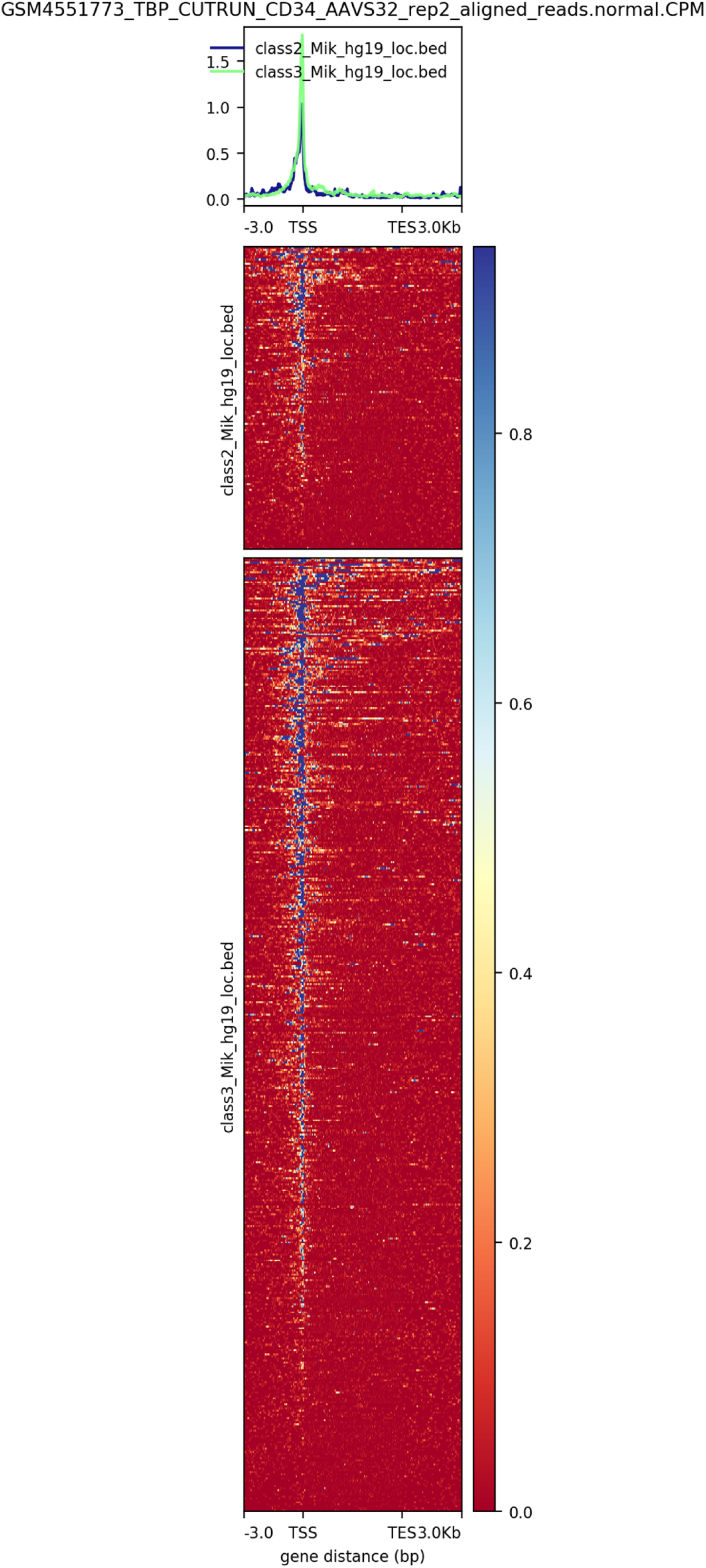
Track plots for TBP occupancy on class II and III genes from the PROseq data. MLLT3 target genes from ChIPseq ^2^ in class II and III in our PROseq data were analyzed for TBP occupancy based on a TBP CUT&RUN study in human CD34 cells ^48^.

## Materials and Methods

### PAR-CLIP

#### Packing of 3xFLAG AF9 lentivirus

Lentiviral particles were packed in 293TN cells in the Abounader and Bushweller Lab at the University of Virginia. Cells were transfected with pCDH 3XFLAG AF9 and viral protein vectors. Following viral packaging supernatant was harvested and used for infection of HEL cells.

#### Infection of Human Erythroleukemia (HEL) cells

HEL cells were infected using filtered supernatant from lentiviral packing experiment above. Briefly, 5 x 10^4^ cells were plated and allowed to incubate for 24 hours. Following incubation cells were infected with FLAG-AF9 lentivirus by spinning cells with 50% (v/v) supernatant and 8 μg/mL polybrene 30 minutes and then allowing them to incubate for 24 hours. Following this media was changed and cells were pooled and grown 48 hours to achieve a higher density to allow for selection.

#### Selection of 3XFLAG-AF9 infected HEL cells

Infected cells from above were incubated with RPMI 1640 supplemented with 10% FBS supplemented with 10 ug/mL Blasticidin. Cells were incubated with Blasticidin for 7 days when control samples were >99% eradicated. Following this, selected cells were harvested and grown to large cell numbers.

#### Crosslinking AF9 RNA binding proteins with 4-Thiouridine

Cells were incubated with 200 µM 4-SU for 4 hours and harvested. Cells were washed with PBS and resuspended in 20 mL PBS. Cells were crosslinked using a Stratagene 1500 Crosslinker for 0.2 J/cm^2^ at 365 nM (Spitzer et al., 2014). Cells were spun down and wet cell pellets flash frozen.

#### Pulldown of FLAG-AF9 and labeling using ^32^P ATP

Cells were lysed using NP-40 buffer and digested with 0.1U RNAse T1. Following this FLAG-AF9 was pulled down using anti-FLAG magnetic beads. Co-precipitated RNA was then labeled with ^32^P using Poly nucleotide kinase (PNK) and ^32^P labeled ATP.

#### SDS gel of FLAG-AF9 and bound RNA

Using the ^32^P labeled samples a SDS-PAGE gel was ran and transferred to a nitrocellulose membrane. This membrane was imaged using a Typhoon 9400 radioactive phosphorimager instrument (GE).

#### PAR-CLIP 3’ adapter ligation of RNA

Using the data from SDS gel imaging, bands corresponding to FLAG-AF9 were excised and digested with Proteinase K to release RNA from the nitrocellulose membrane. RNA was precipitated and resuspended in a ligation mixture to ligate 3’ adapter overnight. Following this overnight incubation samples were run on a 15% polyacrylamide urea gel (UREA systems). Gels were imaged using a Typhoon 9400 radioactive phosphorimager instrument (GE).

#### PAR-CLIP 5’ adapter ligation of RNA

Using the data from 3’ radioactive imaging, bands of the correct size were excised from the polyacrylamide urea gel. Additional larger footprint bands were also excised as well. RNA was eluded from excised bands and precipitated. Samples were resuspended in a 5’ adapter ligation mixture and was incubated for 1 hour at room temperature. Following this overnight incubation samples were run on a 12% polyacrylamide urea gel (UREA systems). Gel were imaged using a Typhoon 9400 radioactive phosphorimager instrument (GE).

#### Preparation and amplification of cDNA library

Using data from the 5’ imaged radioactive gel, bands were excised corresponding to the correct size libraries. This RNA was reverse transcribed using Superscript III Reverse Transcriptase system (Invitrogen). This cDNA library was purified and amplified by PCR for 10 samples. The resulting DNA was purified and run on a Pippen prep gel (Sage Science) to separate correct sized DNA. The resulting DNA was used to run a pilot PCR in which the correct number of cycles to yield the highest cDNA library concentration without amplifying artifact DNA. Following this a full-scale PCR reaction was run using the correct cycle count. Resulting DNA was purified and again run on a Pippen prep (Sage Science). Final library from Pippen prep was examined using Agilent Tape station (Agilent) to obtain size and concentrations of library. Samples were multiplexed and run on a HiSeq2500 sequencer (Illumina).

#### Processing of PAR-CLIP data

Sequenced data was demultiplexed and sorted using cutadapt (M. Martin, 2011). Sorted data was aligned using Tophat alignment software (Trapnell, Pachter, & Salzberg, 2009). Aligned data was processed using PARalyzer software (Corcoran et al., 2011). Output was examined and sorted using Excel.

### Biochemical experiments

#### Expression and purification of AF9 YEATS domain

For biophysical studies AF9 YEATS domain was expressed utilizing a pET22b vector encoding for MLLT3 AA 1-140 (YEATS domain). This vector was transformed into Rosetta 2 (DE2) cells, modified BL21 cells that are T7 lysogens. These cells were used to inoculate a small starter culture of Terrific Broth (TB) for overnight outgrowth. This culture was used to inoculate a larger expression culture comprised of TB supplemented with 0.4% (v/v) Glycerol. This culture was grown to a density of OD_600_= 1.0, following which temperature was reduced to 15°C. When temperature was achieved the culture was supplemented with 1mM IPTG to induce expression for 12-16 hours. Cells were pelleted at 3500 x*g* and either frozen at –80 °C for future use or purified immediately.

Cell pellets were resuspended in 500mM KCl, 25 mM Imidazole, 1 mM DTT, 25 mM Bis Tris pH=6.9 supplemented with Roche cOmplete™ EDTA-free protease inhibitor tablets and 10% (v/v) Glycerol. Utilizing an Emulsifex C3© cell disrupter, cells were lysed at 10,000 psi for three cycles. Lysate was centrifuged for 60 min at 35,000 x*g* to pellet cellular debris. Clarified supernatant was loaded on to a bulk Qiagen® Ni-NTA agarose resin and allowed to pass through by gravity. Proteins were washed on column using a high-salt buffer (1 M KCl, 40 mM Imidazole, 1 mM DTT, 25 mM Bis Tris pH=6.9), after 5 column volumes Pierce™ Universal Nuclease was added to 2 column volumes and allowed to digest 24 hours at 4°C. The following day high salt buffer was passed through the column until OD_260_<0.1. To elute, a high imidazole buffer was employed (600 mM KCl, 300 mM imidazole, 1 mM DTT, 25 mM BisTris pH=6.9), proteins were eluted until OD_260/280_ rose above 0.8 or OD_280_ fell below 0.1.

Following affinity purification, ion-exchange chromatography was employed. AF9 YEATS domain was diluted to a salt concentration of 50 mM KCl and loaded onto a pre-equilibrated SP sepharose column. The column was washed using a low salt buffer (50 mM KCl, 5 mM DTT, BisTris pH=6.5) and gradient eluted using a high salt buffer (1 M KCl, 5 mM DTT, BisTris pH=6.5). Fractions containing AF9 YEATS domain were consolidated and exchanged into a storage buffer (600 mM KCl, 5 mM DTT, BisTris pH=6.5, 10% (v/v) Glycerol) and frozen at –80 °C or into appropriate assay buffer.

#### Expression and purification of ^15^N AF9 YEATS domain

For NMR studies AF9 YEATS domain was expressed similarly to protein for FP studies. However, European Molecular Biology Laboratory (EMBL) media was used for selective ^15^N labeling. Media was supplemented with 1g/L (^15^NH_4_)_2_SO_4_ and 5mL/L ^15^N Bioexpress (Cambridge Isotopes Laboratories). Expression were induced with 1mM IPTG when cell density reached OD_600_=0.8.

Following expression, the protein sample was purified similarly to that used in biophysical studies.

#### NMR measurements and CSP calculations

Backbone resonance assignments for the AF9 YEATS domain were obtained using standard triple resonance techniques suing ^13^C/^15^N-labeled protein. All NMR experiments for AF9 YEATS domain determination of consensus sequence and titration of consensus RNA were conducted with a 50 µM sample of AF9 YEATS in 25 mM Bis-Tris/MES, 100mM NaCl, 1mM DTT, pH 6.0 at 25 degrees Celsius using a Bruker 800 MHz magnet equipped with a cryogenically cooled probe. Experiments were run for 1 hour using 16 scans and 128 points collected in the indirect dimension. All data were collected and processed using NMR pipe (Delaglio et al., 1995). Spectral data was observed using CCPnmr Analysis(Skinner et al., 2016, 2015). CCPnmr was also used to calculate CSPS using the formula(Yuan, Davydova, Conte, Curry, & Matthews, 2002):

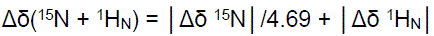

Further calculation of consensus sequence was performed in Excel. While calculation of consensus sequence binding affinity was calculated using CSPs in OriginLabs.

For the interaction study of wt AF9 YEATS and K63E/K67E (dm) with 7SK SL4, protein samples were prepared as previously described. The NMR buffer consisted of 50 mM potassium phosphate, 50 mM KCl, 2 mM DTT, pH 7.0. Proteins and 7SK SL4 were mixed in a 1:1 molar ratio (250 μM each) and incubated at room temperature for 1 hr prior to NMR data acquisition at 298K. ^15^N-^1^H HSQC were recorded for both the free proteins and proteins with 7SK SL4. Each NMR experiment was run for 1.5 hours with 20 scans and 128 points were acquired in the indirect dimension. Chemical shift perturbations (CSPs) were calculated using the following equation and plotted as a function of amino acid residue number using MS Excel.

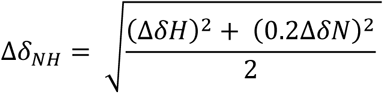

#### Fluorescence Polarization measurement of AF9 Affinity for RNA

For FP assay AF9 YEATS domain purified as described above was used. For all oligos experiments were carried out in 25 mM Bis-Tris/MES (pH 6.0), 100mM NaCl, 1mM DTT buffer. Purified AF9 YEATS protein was titrated into a solution of 20 nM 6-[FAM] conjugated oligos, starting at 10 µM and serially diluted with 3-fold dilutions 8 or 12 times. Samples were incubated for 30 minutes at room temperature. Fluorescence was measured using a PHERAstar plate reader (BMG Labtech) and fluorescence polarization (FP)values were calculated using perpendicular and parallel fluorescence values. FP values were fit to a sigmoidal curve and K_D_ was determined usingGraphpad Prism software.

#### Design and production of AF9 YEATS domain binding mutants

Mutations of AF9 YEATS domain were selected from the 24 residues used in determination of consensus sequence. Primers were designed for mutants selected using PrimerX (https://www.bioinformatics.org/primerx) and ordered from Eurofins Genomics. Quick change reactions were performed on a Biorad thermal cycler using gradient annealing. Following quick change PCR, plasmid was digested with DPNI and transformed into *E. coli* Turbo cells (New England Biolabs). Colonies were sequenced at Eurofins Genomics to confirm mutation.

Mutant proteins were expressed and purified similarly to above described protocol for AF9 YEATS WT construct.

### Hematopoiesis studies

#### Cloning of Double Mutant AF9 lentiviral construct

Mutagenesis reactions were performed by Genscript Biotech Corporation® mutations were incorporated into the same backbones of vectors used to pack virus for HEL viral transduction as well as for Mouse primary cells. Lentiviral constructs were shipped to the lab of Dr. Nancy Zeleznik-Le where differentiation and gene expression analysis experiments were performed.

#### Lentivirus production

293T packaging cells were co-transfected with MLLT3 constructs (*CSIEm-MLLT3 WT-GFP,* for murine lin-BM as describe in ^57^); (pCDH-3xFLAG-HA-MLLT3 WT, pCDH-3xFLAG-HA-MLLT3 K63E K67E for HEL MLLT3 KO cells) and lentiviral packaging constructs (pCMV-dR8.2, and pCMV-VSVG) using the CalPhos Mammalian Transfection Kit (Clontech #631312). Lentivirus containing cell culture supernatants were harvested at 24 and 48hrs. post-transfection, combined, and concentrated 100 fold using Lenti-X Concentrator (Takara #631231) following the manufacturer’s protocol. Virus titers were determined by transduction of 293T cells with concentrated lentivirus followed by GFP^+^ assessment 48hrs. post transduction.

#### Bone marrow progenitor isolation, transduction, and culturing

Bone marrow (BM) was harvested *Mllt3 ^flox/flox^ Rosa26cre^ERT^*^2^ mice. The derivation and functional characterization of these mice will be published separately. Mature BM cells were depleted using the MojoSort™ Mouse Hematopoietic Progenitor Cell Isolation Kit (BioLegend #480003) following the manufacturer’s protocol. The stem and progenitor cell enriched population was cultured overnight at 37C 5% CO_2_ in RPMI-1640+10% FBS+1% Penicillin/Streptomycin (culture media) supplemented with 2-ME [0.05mM], and mSCF, mIL-3, and mIL-6 each at 10ng/mL. The following day 10^5^ cells were transferred to one well of a 24 well ultra-low attachment plate in 300µL culture media containing polybrene [8µg/mL] and CSIEm-*MLLT3 wild type* or CSIEm-*MLLT3 K63E K67E* overexpressing GFP lentivirus. The cells were spinoculated in the lentivirus containing media by centrifugation at 500xg for 45min at 32C. Following spinoculation 300µL of fresh culture media containing 2-ME and cytokines (as above) was added to the well. The cells were then cultured at 37C 5% CO_2_. Cells were harvested from the 24well plate 72hrs after spinoculation. GFP+ transduced cells using a FACSAria (BD) flow cytometer and cultured for 8 days in methylcellulose (StemCell #M3234) containing mFlt3L (10ng/mL), hTPO, hEPO, mSCF, mGMCSF, mIL-3, mIL-6, mIL-11 (each at 20ng/mL) (Peprotech, see supplement), gentamycin (50µg/mL) (Gibco# 15750-060), and 4-hydroxytamoxifen [200nM] (Sigma #T5648) to induce deletion of endogenous *Mllt3*, and subsequently harvested for analysis. Antibody staining to assess hematopoietic cells (see supplement) was performed and cells analyzed with an LSRFortessa flow cytometer (BD Biosciences) and cell surface marker expression assessed using FlowJo (TreeStar Inc., Oregon).

### NES-MLLT3 transformation studies

#### Virus production for NES-MLLT3 and NES-MLLT1 constructs

PLAT-E packaging cells (a gift from Toshio Kitamura)^58^ were used to produce ecotropic retroviruses. The virus-containing medium was harvested 24–48 h after transfection and was used for viral transduction.

#### *Ex vivo* myeloid progenitor transformation assay for NES-MLLT3 and NES-MLLT1 constructs

The myeloid progenitor transformation assay was performed as previously described in detail^45^. Bone marrow cells were harvested from femurs and tibiae of 5-week-old female C57BL/6JJcl (C57BL/6J) mice, from which c-Kit^+^ cells were enriched using magnetic beads conjugated with an anti-c-Kit antibody (Miltenyi Biotec). The c-Kit-positive cells were transduced with a recombinant retrovirus by spinoculation, and plated (4 × 10^4^ cells/sample) in a methylcellulose medium (IMDM, 20% FBS, 1.6% methylcellulose, and 100 µM β-mercaptoethanol) containing murine SCF, IL-3, and GM-CSF (10 ng/mL each). During the first passage, G418 (1 mg/mL) was added to select the transduced cells. Relative *Hoxa9* expression was measured using qRT-PCR after the first passage. The cells were re-plated once every five days with a fresh medium, and colony-forming units per 10^4^ plated cells were quantified during each passage.

#### Fractionation-assisted chromatin immunoprecipitation (fanChIP) for NES-MLLT3 constructs

Chromatin fractions from HEK293T cells were prepared using the fanChIP method as previously described in detail^46^. Cells were suspended in CSK buffer [100 mM NaCl, 10 mM PIPES (pH 6.8), 3 mM MgCl_2_, 1 mM EGTA, 0.3 M sucrose, 0.5% Triton X-100, 5 mM sodium butyrate, 0.5 mM DTT, and protease inhibitor cocktail] and centrifuged (400 × *g* for 5 min at 4 °C) to remove the soluble fraction. The pellet was resuspended in MNase buffer [50 mM Tris-HCl (pH 7.5), 4 mM MgCl_2_, 1 mM CaCl_2_, 0.3 M sucrose, 5 mM sodium butyrate, 0.5 mM DTT, and protease inhibitor cocktail] and treated with MNase at 37 °C for 3–6 min to obtain oligonucleosomes. The MNase reaction was terminated by adding EDTA (pH 8.0) at a final concentration of 20 mM. Equal amounts of lysis buffer (250 mM NaCl, 20 mM sodium phosphate [pH 7.0], 30 mM sodium pyrophosphate, 5 mM EDTA, 10 mM NaF, 0.1% NP-40, 10% glycerol, 1 mM DTT, and EDTA-free protease inhibitor cocktail) were added to increase solubility. The chromatin fraction was cleared by centrifugation (17,700 × *g* for 5 min at 4 °C) and subjected to IP with specific antibodies and protein-G magnetic microbeads (Invitrogen) or anti-FLAG M2 antibody-conjugated beads. The purified materials were then washed five times with washing buffer (a 1:1 mixture of lysis buffer and MNase buffer with 20 mM EDTA) and eluted with elution buffer (1% SDS and 50 mM NaHCO_3_). The eluted material was analyzed by WB.

### Gene expression analysis

#### RNA isolation for RNA-seq

Bone marrow progenitor cells were isolated, transduced, and cultured as indicated above. The cultures were harvested after 8 days. RNA was extracted from mouse bone marrow which was transduced with wild type MLLT3 or MLLT3 K63E K67E mutant followed by deletion of endogenous Mllt3 in the cells using the AllPrep DNA/RNA Mini Kit (Qiagen, #80204), according to manufacturer’s instructions (n=3 per genotype).

#### RNA-seq

Second-stranded libraries with ERCC spike-in controls (ThermoFisher Scientific, #4456740) were prepared by the University of Miami Hussman Institute for Human Genomics Sequencing Core using the TECAN Universal Plus Total RNA-seq Kit with AnyDeplete and 300 ng of RNA as input. Libraries were sequenced on the NovaSeq 6000 with 100 bp paired-end sequencing.

#### RNA-seq alignment

Using Cutadapt (v2.6), adapters were removed from both reads ^59^. Reads were aligned to the mm10 reference genome using the STAR aligner (v2.7.3), specifying the following parameters: STAR –-runThreadN 5 –-genomeDir genome_path –-readFilesIn trim_R1_fullpath trim_R2_fullpath –-outFileNamePrefix name_ –-outFilterType BySJout –outFilterMultimapNmax 20 –-alignSJoverhangMin 8 –-alignSJDBoverhangMin 1 –-outFilterMismatchNmax 999 –-alignIntronMin 20 –-alignIntronMax 1000000 –-alignMatesGapMax 1000000 –-readFilesCommand gunzip –c –-outSAMtype BAM SortedByCoordinate –-outWigType bedGraph –-outWigStrand Stranded –-outWigNorm RPM –-alignEndsType EndToEnd ^60^.

#### Differential gene-expression analysis

Gene counts were calculated using QoRTs (v1.3.0) with the mm10 Gencode annotation file (vM25) that also contained the ERCC spike-ins. Differential gene expression analysis was performed using the package DESeq2 (v1.30.0) with the DESeqDataSetFromHTSeqCount and DESeq functions ^61^. The log-fold change was corrected using the ashr method ^62^. Significant genes were defined as having a fold change >1.5 and p-adjusted <0.05. Regularized log-counts (rld) and the Wald-statistic were generated with DESeq2. For GSEA (v4.0.3) the Wald statistic ranked list was used with the Mouse_Gene_Symbol_Remapping_Human_Orthologs_MSigDB.v7.4.chip chip and the c2.all.v7.4.symbols curated gene set using the weighted enrichment score ^63^.

For the heatmap of genes that are differentially expressed in DM compared to WT, normalized counts were plotted with ComplexHeatmap (v2.6.2) using distance correlation and the average clustering method ^64^.

#### Nuclear Isolation for PRO-seq

Bone marrow progenitor cells were isolated, transduced, and cultured as indicated above. The cultures were harvested after 8 days. Cells were washed in PBS and the nuclei from 2×10^6^ cells were isolated by incubation for 5 minutes on ice in a solution of 10mM Tris, 300mM sucrose, 3mM CaCl_2_, 2mM MgAc_2_, 0.1% Triton-100, and 0.5mM DTT. The lysate was then homogenized with 25 passes using pestle A in a Dounce homogenizer. Nuclei were pelleted and resuspended in a solution of 10mM Tris, 25% glycerol, 5mM MgAc_2_, 0.1mM EDTA, and 5mM DTT, snap frozen on dry ice, and stored at –80C until PRO-seq was performed.

#### PRO-seq

PRO-seq was performed on nuclei extracted from mouse whole bone marrow cells that were transduced with MLLT3 WT or MLLT3 K63E K67E overexpressing lentivirus followed by cell sorting and culture in methylcellulose containing mFlt3L (10ng/mL]) hTPO, hEPO, mSCF, mGMCSF, mIL-3, mIL-6, mIL-11 (each at 20ng/mL)), gentamycin (50µg/mL), and 4-hydroxytamoxifen (200nM) to induce deletion of endogenous Mllt3. Cells were cultured in methylcellulose at 37C 5% CO2 for 8 days before being harvested for PRO-seq. PRO-seq experiments were performed as previously described^47,65^. Nuclear run-on assays were performed on one million nuclei for 3 minutes at 30 °C using 25μM Biotin-11-ATP/UTP/CTP/GTP (PerkinElmer). Total RNA was extracted, and fragmentation was performed with 0.2M NaOH for 10 min on ice. Biotinylated nascent RNA was purified by streptavidin beads M-280 (Invitrogen, 11205D), and the 5’ cap was removed using RppH (NEB, M0356S). T4 polynucleotide kinase (NEB, M0201) was then used to repair the 5’ hydroxyl terminus. Following adapter ligation, SuperScript III (Thermo Fisher Scientific, 18080044) was used for reverse transcription. After PCR amplification, 140–350 bp libraries were size-selected by AMpure XP (Beckman Coulter) and sequenced on a NovaSeq 6000 (Illumina) with single-read runs.

#### PRO-seq analysis

After sequencing, trimming of raw fastq data, genome alignment and spike-in normalization were performed as previously described ^47,65^. Briefly PRO-seq reads were trimmed by Cutadapt 1.14^66^ and Trimmomatic v0.32 ^67^ and aligned to the mouse genome (mm10) or the drosophila genome (dm3) by bowtie 1.1.2 ^68^ PRO-seq signal was normalized by the number of reads mapped to spike-in dm3 genome, and then converted to bigwig data, which were used for downstream analyses. To determine the accurate TSS for each gene, we separated the mapped reads from PRO-seq by strand and individually call peaks using MACS2.1.2 ^69^ with the following parameters: –-nomodel, –-extsize 200 and –narrow to identify narrow peaks. We merged the common peaks from each sample (WT and DM) for each stranded peak and used BedTools v2.28 ^70^ command closest to annotate the peaks against mouse Gencode (v25). Peaks with distance >100nt from an annotated TSS region were discarded to avoid multiple assignments to the same promoter. When multiple TSS from the same reference were detected in the same PRO-seq peak, we selected the most upstream isoform for analysis. Also, when a gene has multiple isoforms, the distance between these TSS was > 350nt and they were assigned to different PRO-seq peaks, we kept all isoforms for analysis. Next, we calculated the RPKM values in promoter region (–50 nt from TSS to + 300 nt) and the region corresponding to the gene body (+301 nt from TSS to TES) for all transcripts used in the analysis (n = 13,278). These values were used as input for the traveling matrix (TM). The TM can be visualized as a 2D distribution by using the log_2_ ratio (RPKM KO over RPKM WT) of the promoter region (X axis) and gene body (Y axis). The relative difference generated by this 2D representation can be categorized in four different groups: Class 1: log_2_ promoter > 0 and log_2_ gene body < 0; Class 2: log_2_ promoter < 0 and log2 gene body < 0; Class 3: log_2_ promoter > 0 and log2 gene body > 0 and Class 4: log_2_ promoter < 0 and log2 gene body > 0.0 The read counts in the gene body region were used to calculate and determine the differential gene expression (DEG) between conditions using DESeq2 ^61^. Differentially expressed genes were identified by using a 1.25-fold change cutoff (FDR<0.05) and applying a minimum gene expression filter for the final set of differentially expressed protein-coding genes (0.3 RPKM in both replicates in at least one of the conditions). For GSEA (v4.0.3) the Wald statistic ranked list was used with the Mouse_Gene_Symbol_Remapping_Human_Orthologs_MSigDB.v7.4.chip chip and the c2.all.v7.4.symbols curated gene set using the weighted enrichment score ^63^. For comparison to human HSPC MLLT3 targets and TBP binding sites, gene names from Class II and Class III genes were converted to human gene symbols and for those that were also direct MLLT3 targets in human hematopoietic stem and progenitors (HSPC) ^2^, the TBP signal (counts per million) in human HSPC ^71^ was plotted over the corresponding genebodies using deeptools.

#### MLLT3KO HEL transduction and selection

HEL cells with MLLT3 knocked out using CRISPR/Cas9 technology were generated by Abcam. Homozygous knockout was confirmed via sequencing. MLLT3KO HEL cells were cultured in RPMI 1640+10% FBS+primocin [100ug/mL] (Invivogen #ANT-PM-05). On the day of transduction 10^5^ MLLT3 KO HEL cells were transferred into one well of a 24 well ultra low attachment plate in 500uL culture media containing polybrene [8µg/mL] and HA tagged *MLLT3 wild type* or *MLLT3 K63E K67E* overexpressing lentivirus. The cells were spinoculated in the lentivirus containing media by centrifugation at 500xg for 45min at 33C. Following spinoculation 500µL of fresh culture media was added to the well. The cells were then cultured at 37C 5% CO_2_. 72hrs after transduction the cells were harvested, pelleted, and resuspended in culture media containing 10µg/mL blasticidin to select transduced cells. The cells were cultured at 37C 5% CO_2_ under blasticidin selection for 11 days at which point the blasticidin was reduced to 5µg/mL and the cells were expanded to sufficient numbers for subsequent experiments.

#### MLLT3 HA-ChIP

MLLT3 (WT or K63E K67E) overexpressing MLLT3KO HEL cells were cultured as above, harvested, crosslinked in 1.1% formaldehyde for 10min. at room temperature, quenched with glycine, washed with PBS, and pelleted. Chromatin was prepared using the iDeal ChIP-qPCR kit (Diagenode #C01010180) following the manufacturer’s protocol. Harvested chromatin was sheared with eight cycles (30s on, 30s off) at 4C using a Bioruptor Pico (Diagenode). Chromatin bound HA tagged proteins were immunoprecipitated with an anti-HA antibody (Cell Signaling Tech #3724S), eluted, and purified using the iDeal ChIP qPCR kit (Diagenode) protocol. Immunoprecipitated chromatin was interrogated using primers located near the transcription start sites of various genes. Primers were designed using NCBI Primer BLAST. SYBRgreen qPCR was performed using SsoAdvanced Universal SYBR Green Supermix (Bio-Rad# 1725271) on a QuantStudio 6 PCR system (Applied Biosystems).

### Bioinformatic analysis to compare overlap of 7SK and MLLT3 binding sites for class II and III genes from the PRO-Seq

#### Data Preparation

The Mouse PROseq data with genes from class2 and class3 were compared with the Mouse ChIRP-seq 7SK data downloaded from GEO (GSE69141: GSE69141_mm9_7SK_ChIRP-seq_mES_WT.bw). The ChIRP-seq 7SK data was first converted from bw to bedGraph file format and then converted to mm10 co-ordinates using liftOver ^72^ UCSC utilities program as it was originally aligned to mm9 genome build. After conversion to mm10 coordinates, there were a series of steps done to remove regions with zero coverage, sorted them and merged the consecutive regions using bedtools merge ^70^ program with a maximum distance (–d) of 100 allowed between the regions. The length distribution of the 7SK ChIRP-seq was then observed and since majority of regions were of length between 140 to 150, we considered the regions that are greater than 100 in length.

#### Comparison of PROseq data with Mouse ChiRP-seq (GSE69141)

The regions from the PROseq class2 and class3 genes were individually compared with the above prepared Mouse ChIRP-seq data using bedtools intersect ^70^ program.

#### Comparison of Human 7SK ChIRP-seq (GSE69141) with Human MLLT3 ChIP-seq data (GSM3738886): Data Preparation

Both ChIRP-seq (GSE69141_hg19_7SK_ChIRP-seq_HeLa.bw) and ChIP-seq data (GSM3738886_K562_1_MLLT3_Gt.bw) belongs to the hg19 and were first converted from bw to wig format bigWigToWig UCSC tools ^72^. After conversion, there were a series of steps done to remove regions with zero coverage, sorted them and merged the consecutive regions using bedtools merge ^70^ program with a maximum distance (–d) of 100 allowed between the regions. Then based on the length distribution of the 7SK ChIRP-seq we considered the regions that are greater than 100 in length for both ChIP-seq as well as ChIRP-seq data.

#### Comparison of 7SK ChIRP-seq (GSE69141) with Human MLLT3 ChIP-seq data (GSM3738886)

The regions from the 7SK ChIRP-seq and MLLT3 ChIP-seq were compared using bedtools intersect ^70^ program. The overlapped regions were then annotated using ChIPseeker::annotatePeak ^73^ function in R.

#### Identifying common genes between ChIRP-seq_CHIP-seq_Overlap genes and the ChIRP-seq_PROseq_Overlap genes

The annotated genes from the overlapped regions between the Human 7SK ChIRP-seq and Human MLLT3 ChIP-seq were compared with the Mouse 7SK ChIRP-seq and Mosue PROseq overlapped genes. First the homologous genes symbols in human were identified for the mouse 7SK ChIRP-seq and mouse PROseq overlapped genes using the BioMart (https://useast.ensembl.org/info/data/biomart/index.html) program from Ensembl. The overlap of genes between the two groups were performed in R using intersect function.

